# Hemilineage-specific expression of the RHG genes *Grim* and *Reaper* sculpt neural network composition during *Drosophila* neurogenesis

**DOI:** 10.1101/2024.02.11.579841

**Authors:** Connor J Sproston, Julia E Rak, Shu Kondo, Darren W Williams

## Abstract

During development, nervous systems generate an incredible diversity of different cell types. One mechanism that appears critical for controlling whether a specific subtype is present or absent, and ensuring that neuronal numbers are appropriate for specific regions, is programmed cell death.

Here we show that the precisely patterned hemilineage-based cell death found in the ventral nerve cord of *Drosophila melanogaster* is controlled by the transcription of the pro-apoptotic RHG genes *reaper* and *grim*.

Using smFISH we show that *reaper* and *grim*, but not *hid*, are expressed in postembryonic lineages during neurogenesis. By generating novel T2A-GAL4 knock-in tools we have been able to map *reaper* and *grim* expression on a lineage-by-lineage basis and show diverse expression patterns within different hemilineage populations. Analysis of null mutants for both genes reveal a ‘division of labour’ with distinct combinations being important in sculpting the nervous system.

Investigation of a specific sexually dimorphic lineage reveals temporal patterning of hemilineage-specific programmed cell death through Grim and Reaper function underpins the development of sex-specific circuitry in the VNC.

Our findings suggest that the precisely patterned expression of these pro-apoptotic genes is a critical fate determinant that generates segmental-specific circuity and instructs the appropriate assembly of sexually dimorphic neural circuits in the VNC.

## Introduction

Nervous systems contain an incredible diversity of cell types that are intricately organised into complex functional networks (Simon and Konstantinides, 2021). These networks arise through the integration of various developmental processes that ensure the generation of the appropriate number of specific cell types in the right location, that then elaborate ordered patterns of synaptic connectivity. One process that appears to be universally deployed during the construction of nervous systems is programmed cell death (PCD) by apoptosis.

PCD is seen extensively during the development of the central nervous system of; nematode worms (Sulston and Horvitz., 1977, Ellis and Horvitz., 1986), lower vertebrates such as fish (Biehlmaier *et al*., 2001, Williams *et al*., 2000), frogs (Lamborghini., 1987, Lamb *et al*., 1989, Yeo and Gautier., 2003), as well as birds (Homma *et al*., 1994, Hamburger and Levi-Montalcini., 1949, Oppenheim, 1991) and mammals (Ikonomidou *et al*., 1999, Blanquie *et al*., 2017, Wong *et al*., 2018).

In *Drosophila*, a highly stereotyped pattern of neuronal death is seen in the postembryonic development of the brain and the ventral nerve cord (VNC) (Truman *et al*., 2010, Lin *et al*., 2010, Kumar *et al*., 2009 Bertet *et al*., 2014, Lovick *et al*., 2017, Pop *et al*., 2020). This patterned PCD is observed in the lineages of Type-I Neurobalsts (NBs) of the VNC and Brain that divide in a self-renewing fashion to generate ganglion mother cells (GMCs) which undergo terminal division to generate a Notch-ON ‘A’ daughter and a Notch-OFF ‘B’ daughter (Spana and Doe, 1996, Lundell *et al*., 2003). Each Type-I NB thus has the potential to produce a lineage of daughter cells comprised of two half-lineages or hemilineages, an A hemilineage and a B hemilineage (Fig.1A). During postembryonic development of the larval ventral nerve cord patterned developmental neuron/cell death is observed to stereotypically remove specific and identifiable thoracic hemilineage populations born from these NBs (Truman *et al*., 2010) (Fig.1B). This patterned death is extensive and occurs throughout the whole neurogenic period, we term this, ‘hemilineage-specific programmed cell death’ (Pop *et al.,* 2020).

**Figure 1:**
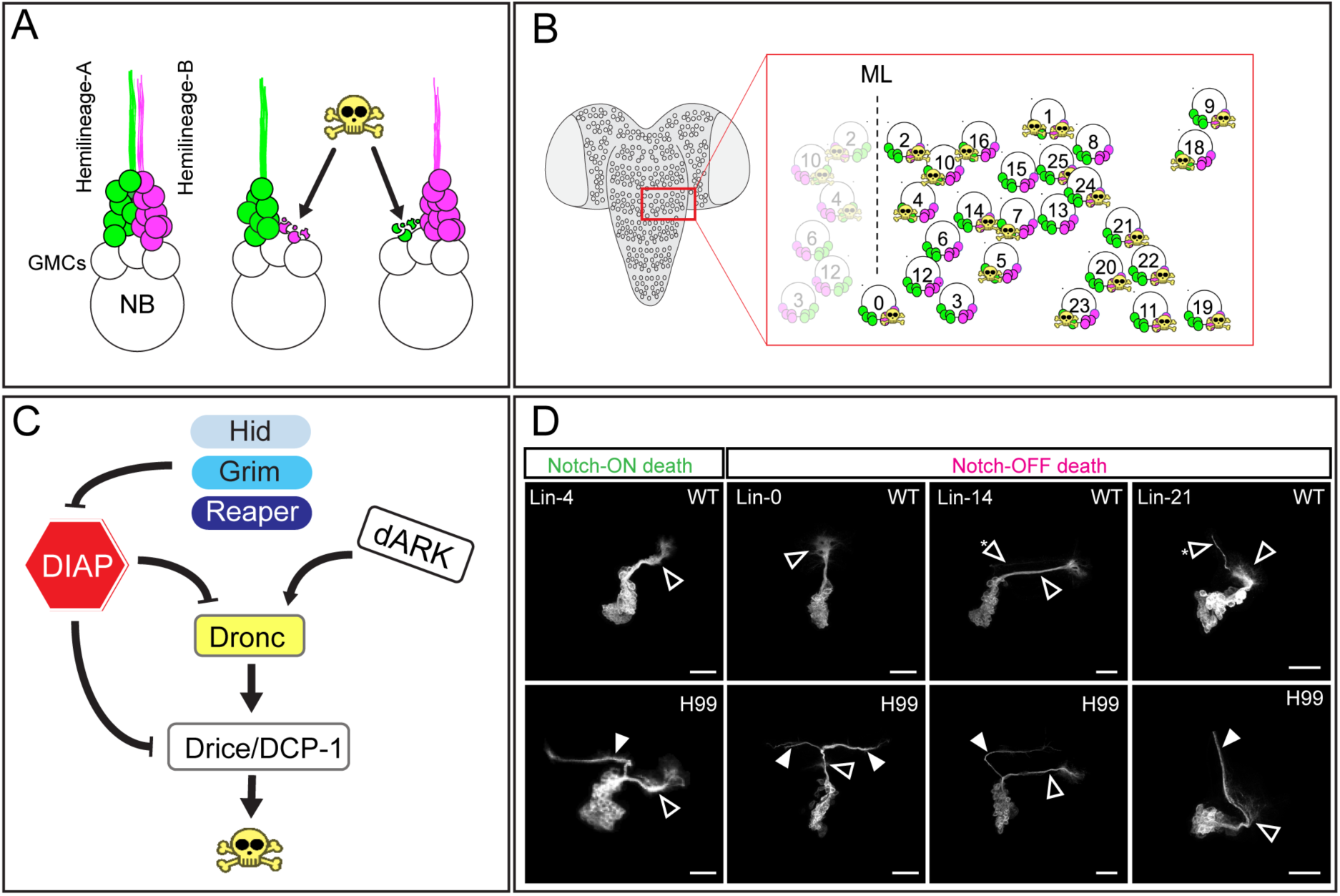
(**A**) **N**euro**b**lasts (NB) divide to give rise to **G**anglion **M**other **C**ells (GMC’s) which divide terminally to generate an **A** (Notch-ON) and **B** (Notch-OFF) daughter. This results in a lineage of postmitotic neurons which is further subdivided into an A hemilineage and B hemilineage. In lineages where hemilineage **P**rogrammed **C**ell **D**eath (PCD) occurs either the A or B cells are removed through apoptosis soon after their birth. This generates ‘monotypic’ lineages made up largely of a single hemilineage. (**B**) The pattern of death within the hemisegmental array of neuroblast lineages and their hemilineages is stereotypical with 18/26 NB lineages displaying hemilineage specific PCD (**C**) Schematic displaying the core aspects of the pro-apoptotic signaling cascade in *D.melanogaster,* the RHG proteins, Reaper, Hid and Grim, the key regulators of DIAP1 (*Drosophila* Inhibitor of Apoptosis). DIAP1 the inhibitor of the initiator caspase Dronc (caspase-9 homolog) and the effector caspases Drice and DCP-1 (caspase-3 homologs). Drice and DCP-1 when active cleave cellular proteins and result in death. Also shown is the initiator caspase co-factor dARK (*Drosophila* Apaf-related killer). (**D**) Clonal removal of the RHG genes **R**eaper, **H**id and **G**rim in postembryonic NB lineages that feature PCD in Notch-ON hemilineage-A (Lineage-4) and Notch-OFF hemilineage-B (Lineage-0, 14, 21) results in the rescue of the normally doomed hemilineage (filled arrowhead). ‘wt ‘non-doomed hemilineage as shown with open arrowheads. Asterisk (*) depicts examples where small number of neurons from ‘doomed ‘hemilineage are observed in wt conditions (Lineage 14 and 21) these populations are show expansion when PCD is blocked.

The molecular and behavioural interrogation of the surviving hemilineages shows that each specific hemilineage population shares core molecular identities, expresses specific neurotransmitter repertoires and elicits specific behavioural outputs upon activation (Lacin *et al*., 2019, Harris *et al*., 2015). This suggests that the hemilineage units are fundamental functional units of the adult VNC. It is likely that the differential deployment of patterned PCD in the VNC of different species not only ensures that only the ‘correct’ hemilineages are generated during development of *Drosophila*, but may be one way in which evolutionary differences in the hemilineage composition of the CNS may emerge in insects (Pop *et al*., 2020).

Despite its importance during the assembly of the adult network of flies, how hemilineage-specific cell death is orchestrated with such precision is currently unknown. In *Drosophila* the pro-apoptotic RHG (Reaper, Hid, Grim) genes play a key role in regulating cell death during development by disinhibiting IAPs/inhibitors of apoptosis (Fig.1C) (Goyal *et al*., 2000, Lisi *et al*., 2000, Yoo *et al*., 2002). Various studies have proposed the model that RHGs have complex patterns of regulation and transcription during embryonic and post-embryonic periods of development. The regulation of both *reaper* and *grim* appears to underpin a number of Hox mediated neuronal deaths in the embryo (Miguel-Aliaga and Thor., 2004, Rogulja-Ortmann *et al*., 2008) and a large wave of neuroblast death that specifically targets the abdominal neuroblasts during postembryonic neurogenesis (Peterson *et al.,* 2002 Bello *et al.,* 2003). Alongside the regulation of adult neuronal numbers by neuroblast death, steroid hormone-gated neuron death appears to play a key role during metamorphic remodelling (Winbush and Weeks, 2011) and then in the adult during the removal of neurons used for eclosion and wing-spreading behaviour (Robinow *et al*., 1993, Kimura and Truman., 1990).

Within the developing adult VNC Reaper and Grim are required for the death of the postembryonic octopaminergic neurons generated by the median neuroblast lineage MNB lineage 0 (Pop et al., 2020). Whether this is the case for other hemilineages in the VNC has remained unknown. Here we uncover how the spatial expression patterns the RHG genes *hid, grim, reaper* and *sickle* map onto the lineages of the developing postembryonic VNC and how these pro-apoptotic genes might function to regulate the highly stereotypical patterns of PCD we have previously reported (Truman et al., 2010). We show that *reaper* and *grim*, are the only RHG genes transcribed and translated within the developing VNC neuronal lineages. Furthermore, we show that *reaper* and *grim* are required for hemilineage specific cell deaths, that there are differential requirements of both within different lineages, and that they regulate the proper development of sexually dimorphic circuits in the VNC. As hemilineage-specific PCD is known to be critical for generating segment-specific circuit motifs within and between species this work paves the way for understanding the molecular regulation of this fundamental developmental mechanism.

## Results

### Hemilineage-specific PCD within the developing postembryonic thoracic nervous system requires pro-apoptotic RHG proteins

The three pro-apoptotic genes *reaper*, *hid* and *grim* (RHG) play a major role in gating developmental and physiological cell deaths in *Drosophila* (White et al., 1994, 1996; Grether et al., 1995; Chen et al., 1996) (Fig.1C). MARCM clonal analysis using the H99 deficiency, which deletes *reaper*, *hid* and *grim* (White *et al*., 1994), allows us to generate NB lineage clones that lack the RHG genes in an otherwise wild-type background and analyse the composition of hemilineage populations in the absence of an RHG-mediated apoptotic drive.

Analysis of thoracic lineages, which display hemilineage-specific PCD under normal conditions, suggest that the clonal removal of the RHG genes is sufficient to block death of the doomed hemilineage component. When we generate H99 clones of lineages where only a single hemilineage is normally produced, such as lineages 0, 4, 14 and 21, we observe additional ectopic bundles along with the normally present wild-type neurite bundle (Fig.1D). Critically, these ectopic neurite bundles match the anatomy and trajectory of ‘doomed’ hemilineages observed after the removal of the initiator caspase Dronc (Truman *et al*., 2010). Lineages where hemilineage A (Notch-ON) undergoes PCD (Lin-4 Fig.1D) and those where the hemilineage B is targeted (Notch-OFF) (Lin-0, 14, 21 Fig.1D) are equally impacted when all three RHGs proteins are removed with the *H99* deficiency.

These data suggest that some combination of the three apoptotic genes, *reaper*, *hid* and *grim* covered by *H99* is likely required within postembryonic lineages for them to undergo hemilineage-specific programmed cell death (Fig.1D). This prompted us to investigate the expression and individual requirement of these genes within the developing larval thoracic nervous system to understand the wider role they play during this developmental patterned PCD (Pop *et al*., 2020).

### Hid, Grim and Reaper are expressed within the VNC during larval life

To explore the individual, tissue level expression patterns of *reaper*, *hid* and *grim* in the thoracic VNC we designed single molecule fluorescence *in situ* hybridisation (smFISH) probe-sets for each gene (Fig.2B-D) and probed 3^rd^ larval instar (L3) nervous systems. During this stage we know hemilineage-based PCD is widespread within the neurogenic zone a cortex of cells that contains neuroblasts and newly born neurons (Fig.2A, A’). Each of the RHG genes displayed a distinct pattern of expression within the developing thoracic VNC. Our *hid* specific probe-set consistently shows transcripts within a population of midline glia at wandering L3. These cells occupy the dorsal midline of the VNC, and sit outside the neurogenic zone where we see no clear evidence *hid* mRNA foci above background (Fig2B-B’). To ensure that the probe-set is recognizing *bona fide* Hid transcripts we ectopically expressed *hid* using an inducible *hs-hid* transgene and were able to see robust smFISH signal throughout the whole VNC (Supp. Fig.1).

**Figure 2:**
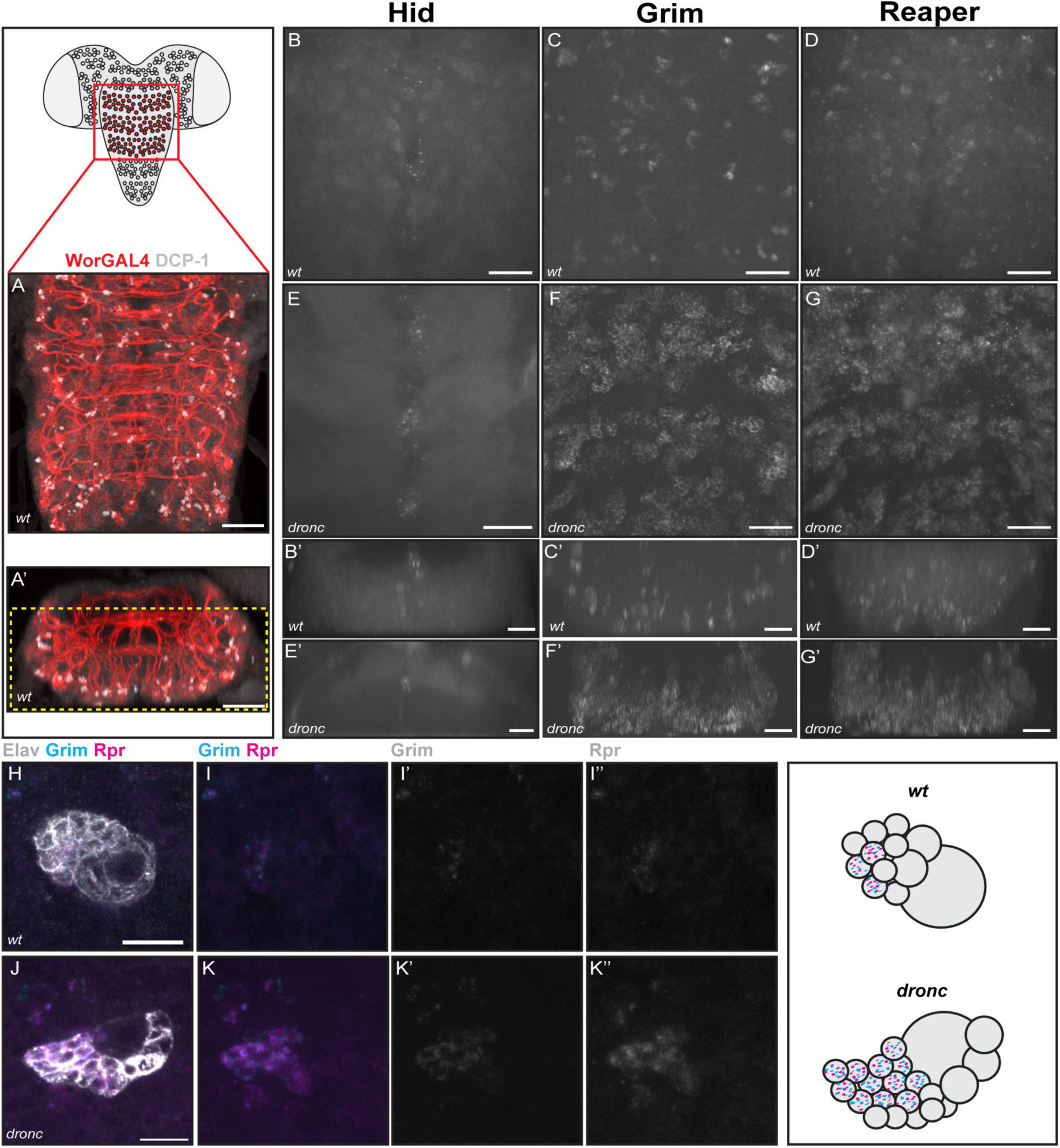
(**A-A’**) *Worniu-GAL4* and staining for the active form of the effector caspase DCP-1 reveals the distribution of hemilineage programmed cell death within the thoracic ventral nerve cord (VNC) during the 3^rd^ larval instar (n= 7). Scale bar, 20μm (**B-D**) Maximum intensity projection of the thoracic neuromeres of *wt* animals showing the expression of Hid (**B**, n=3), Grim (**C**, n=3) and Reaper (**D**, n=3) mRNA using smFISH. Scale bar, 20μm. (**B’-C’**) Maximum intensity projection of transverse sections through the thoracic neuromeres of *wt* animals in **B-D** showing the expression of Hid (**B’**, n=3), Grim (**C’**, n=3) and Reaper (**D’**, n=3) mRNA along the D-V axis. Scale bar, 20μm. (**E-G**) Maximum intensity projection of the thoracic neuromeres of *dronc* animals showing the expression of Hid (**E**, n=2), Grim (**F**, n=7) and Reaper (**G**, n=7) mRNA using smFISH. Scale bar, 20μm. (**E’-G’**) Maximum intensity projection of transverse sections through the thoracic neuromeres of *dronc* animals showing the expression of Hid (**E’**, n=2), Grim (**F’**, n=7) and Reaper (**G’**, n=7) mRNA using smFISH. Scale bar, 20μm. (**H,** n=3) *wt Elav-GAL4* MARCM clone showing expression of *Grim* (**I, I’**) and Reaper (**I, I’’**) mRNA within a single lineage, restricted to a small number of putative ‘doomed ‘post-mitotic cells. Scale bar, 10μm. Schematic cartoon to right. (**J,** n=2) *dronc ELav-GAL4* MARCM clone showing expression of Grim (**K, K’**) and Reaper (**K, K’’**) mRNA within a single lineage where hemilineage PCD is blocked showing expansion of domain of Grim/Reaper mRNA expression as compared to *wt* lineages. Scale bar, 20μm. Schematic cartoon to right.

The *grim* probe-set reveals robust transcript signal in clusters throughout the thoracic VNC (Fig2C-C’). These focal accumulations were distributed widely within the neurogenic zone of the VNC, in locations similar to those seen for dying neurons labelled by active-DCP-1 (Fig2A-A’). Similarly, the probe-set targeting *reaper* mRNA show a similar distribution of expression to *grim*, with accumulations of *reaper* mRNA in clusters distributed throughout the whole neurogenic region of the thoracic VNC (Fig2D-D’).

We also generated a probe-set for the fourth member of the family, as *sickle* even though it has not been shown to block death in other tissues it is expressed within the VNC during embryogenesis (Wing *et al*., 2002, Christich A *et al*., 2002, Srinivasula *et al*., 2002). We see no signal within the larval CNS but were able to recapitulate the published expression pattern in embryos suggesting the probes work well (Supp. Fig. 2).

If members of the RHG gene family are driving hemilineage-specific cell death then removing downstream caspase function, preventing the completion of apoptosis, should result in the accumulation/enrichment of ‘rescued’ hemilineages expressing RHGs. Analysis of thoracic nervous systems in animals homozygous for the initiator caspase Dronc reveal a significant increase in smFISH signals for both *grim* and *reaper* (Fig. 2F, F’ and G, G’). It appears that newly born doomed neurons express *grim* and *reaper* and maintain this expression if unable to complete PCD. In contrast, we found no enrichment of *hid* signal within the rescued postembryonic lineages (Fig. 2E-E’), further suggesting that *hid* is not involved in hemilineage-specific cell death. These *Dronc* mutant VNCs also showed no obvious increase in the low level *hid* smFISH signals seen with the probe-set in midline glia.

To ensure that the observed increase in *reaper* and *grim* signal in the *Dronc* null CNS (Fig. 2F, F’ and G, G’) was due to the accumulation of undead cells and not the result of some ‘non-specific’ global induction of these proapoptotic genes, we looked at *reaper* and *grim* transcripts in MARCM clones. In wild-type *elav-GAL4* clones of single bundle lineages i.e. those that undergo hemilineage-specific death, we observe a small cluster of neurons robustly expressing *reaper* and *grim* directly adjacent to a larger cluster of neurons with no measurable signal (Fig2H-I’’). Such small *reaper* and *grim* positive clusters are similar size to the foci we see in wildtype VNCs (Fig.2 C&D). In *Dronc* null neuroblast clones of equivalent lineages, we find an increased number of *reaper* and *grim* expressing cells adjacent to non-RHG-expressing neurons (Fig.2J, K’’). These data suggest that a failure to complete PCD by removing *Dronc* within a single lineage, results in an increase in the number of *reaper* and *grim* expressing cells (Fig.2J, K’’) as was seen in the whole animal mutant CNS.

To summarise, *reaper*, *hid* and *grim* are all expressed within the thoracic VNC of third instar larvae. Our smFISH probes reveal that Hid is not expressed in the neurogenic regions containing postembryonic neuronal lineages, but rather in a midline population of glial cells. In contrast, both *reaper* and *grim* are expressed in small clusters of cells throughout the neurogenic region where hemilineage-specific PCD is known to be widespread. *Dronc* mutants and clones of *Dronc* reveal an enrichment in high *grim/reaper* expressing cells within postembryonic lineage populations that continue to express both genes. Taken together these data appear to support the idea that *reaper* and *grim* are the primary RHG genes driving hemilineage-specific programmed cell death in thoracic VNC lineages.

### Revealing lineage-specific patterns of Grim and Reaper expression within postembryonic neurons using T2A-GAL4 knock-in reporter alleles

To consolidate our *in-situ* based expression data and reveal lineage-specific patterns of RHG expression within doomed hemilineages we generated T2A-GAL4 knock-in alleles for *reaper*, *hid* and *grim* using CRISPR/Cas9 mediated genome editing (Fig. 3A). These T2A-based tools allow us to visualise the transcriptional output of each of the RHGs in detail, as every time the gene is transcribed and translated a functional GAL4 molecule is generated. When used alone with UAS-reporters these tools show patterns of RHG expression within the CNS of third larval instar. The expression patterns observed are distinct for each RHG gene and suggest that each may be subject to unique spatiotemporal transcriptional control (Fig.3B-D). The T2As also highlight the sensitivity of the system revealing ‘sub-lethal’ RHG expression. Importantly, the knock-in for *hid* shows expression in the same cells in which we observed low-level smFISH signals (Fig.2B-B’ and Fig.3B). The fact that these cells do not show morphological hallmarks of PCD may suggest that the mRNA and GAL4 expression we have observed in these midline glia is related to non-apoptotic Hid function (Huh *et al.,*2004). When looking at the expression of *reaper* and *grim* GAL4s we were unable to see expression within the recently born dying postembryonic thoracic neurons and this is likely to be a result of the rapid cell death and clearance prior to measureable reporter expression.

**Figure 3:**
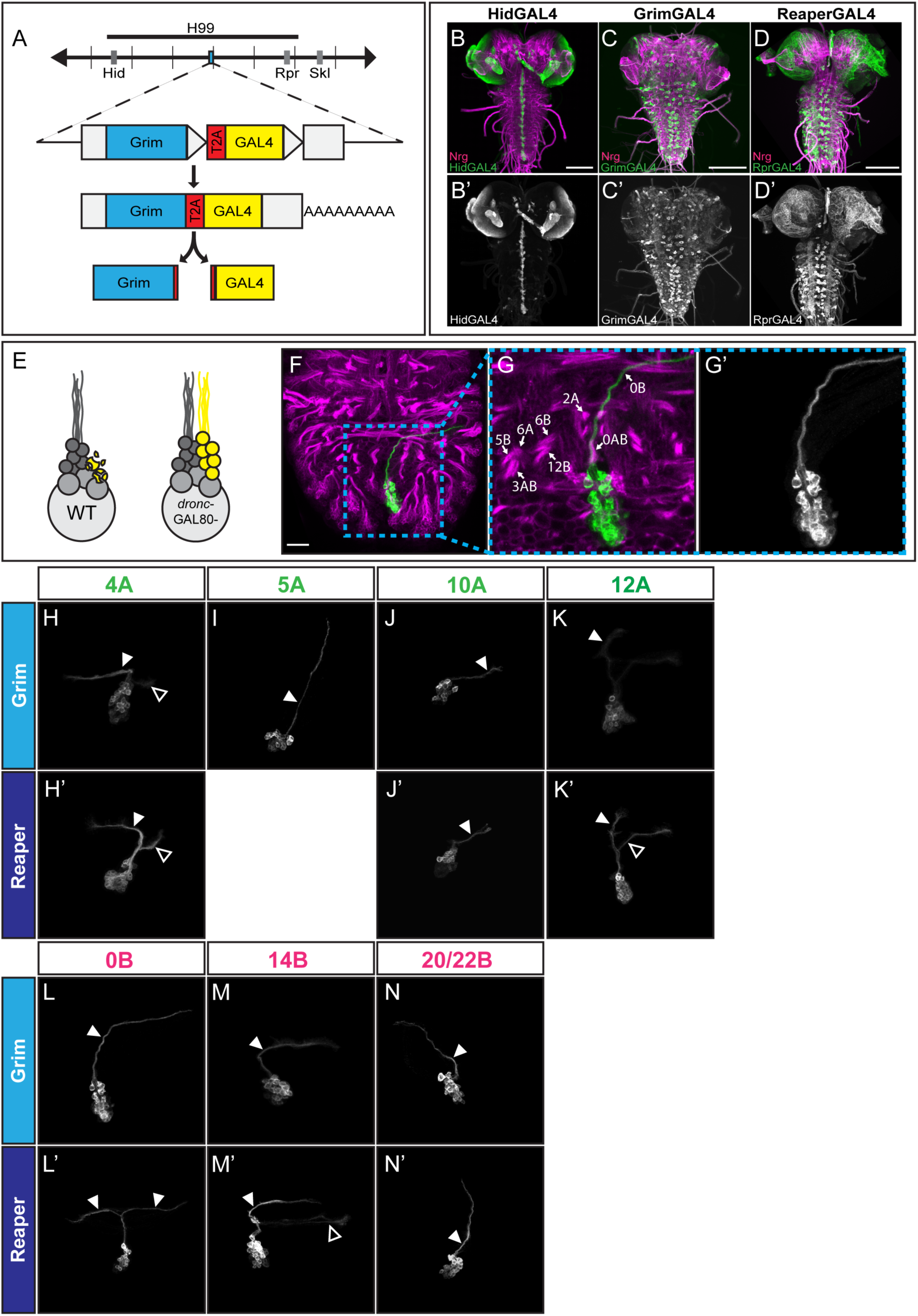
(**A**) Schematic showing the design and structure of T2A-GAL4 RHG gene reporters. (**B-B ‘**n=5) *Hid-T2A-GAL4* expression in the CNS at wL3. (**C-C ‘**n=3) *Grim-T2A-GAL4* expression in the CNS at wL3. (**D-D ‘**n=4). *Reaper-T2A-GAL4* expression in the CNS at wL3. Scale bar, 100μm. (**E**) Schematic representation of RHG-T2A-GAL4 expression in the context of *dronc* MARCM clones. (**F-G’’**) Example hemilineage identification (hemilineage-0B) for *dronc Grim-T2A-GAL4* MARCM clone. Scale bar, 20μm. (**H-H’**) Grim-T2A-GAL4 (**H**, n=18) and Reaper-T2A-GAL4 (**H’**, n=3) expression within hemilineage 4A. (**I**, n=10) Grim-T2A-GAL4 expression within hemilineage 5A. (**J-J’**) Grim-T2A-GAL4 (**J**, n=27) and Reaper-T2A-GAL4 (**J’**, n=4) expression within hemilineage 10A. (**K-K’**) Grim-T2A-GAL4 (**K**, n=10) and Reaper-T2A-GAL4 (**K’**, n=2) expression in hemilineage 12A in T3. (**L-L’**) Grim-T2A-GAL4 (**L**, n=14) and Reaper-T2A-GAL4 (**L’**, n= 3) expression in hemilineage 0B. (**M-M’**) Grim-T2A-GAL4 (**M**, n=19) and Reaper-T2A-GAL4 (**M’**, n=1) expression in hemilineage 14B. (**N-N’**) Grim-T2A-GAL4 (**N**, n=15) and Reaper-T2A-GAL4 (**N’**, n=5) expression in hemilineage 20/22B. Arrow heads show location of rescues ‘doomed ‘ hemilineage projections labelled via their Grim/Reaper expression. Open arrowheads show location of ‘non-doomed’ hemilineages labelled by low level Grim/Reaper expression.

To use these tools to map lineage-specific expression patterns of the RHGs in doomed hemilineages we took advantage of our observation that ‘undead’ *Dronc* null neurons continue to transcribe *reaper* and *grim* (Fig 2D-F). By recombining each T2A-GAL4s onto an FRT, *dronc^null^* chromosome we could use MARCM analysis to reveal hemilineage expression of each RHG through the generation of sparse heat-shock induced lineage clones during neurogenesis. This MARCM-based approach allows detailed characterisation of the expression of the individual RHGs within specific lineages, where all other T2A-GAL4 expression is inhibited by a ubiquitously expressed GAL80. Neuroblast clones for each ‘doomed’ hemilineage population can be readily identified based upon their stereotypical cell body within the array and the trajectory of their neurite bundles within our standardized Neuroglian scaffold (an anatomical scaffold of all primary neurite projections of thoracic postembryonic neurons, revealed by Neuroglian immunohistochemical staining (Fig.3E-F). For example, in a MARCM neuroblast clone for lineage 0, a population previously studied in detail (Pop *et al*., 2020), the primary neurites of the A and B hemilineages emerge from the Lin-0 cell bodies as part of an unpaired (non-bilaterally-symmetrical) lineage located at the posterior of each thoracic neuromere (Fig.3F). The identity of the dorsal projection of hemilineage-0B cells in the lineage clone can be confirmed with reference to the neighbouring bundles in the Neuroglian scaffold (Fig.3G-G’). As observed using our *hid* specific smFISH probe-set, using our *hid-T2A-GAL4* reporter allele we were not able to recover any positively labelled MARCM lineage clones in the VNC and only see clones within the developing optic lobe (Supp. Fig. 3) confirming Hid plays no role in hemilineage PCD in the VNC.

A specific question we wanted to address was whether there may be a differential deployment of Reaper or Grim between A-hemilineages and B-hemilineages i.e. with death in neurons belonging to one hemilineage class always being mediated by Reaper and the other by Grim. In lineages where PCD targets hemilineage-A neurons specifically, we see that the majority of ‘doomed’ hemilineage screened expressed both *reaper* and *grim* (Fig.3H-K’). Hemilineage 4A (Fig.3H and H’) and hemilineage 10A (Fig.3J and J’) display expression of both *reaper* and *grim* within the rescued doomed hemilineage population. Hemilineage 12A represents an example of a lineage where segment-specific hemilineage PCD takes place during larval stages, occurring in hemilineage A population in the third thoracic neuromere. Like the non-segment-specific hemilineage 4A and 10A, hemilineage 12A displays robust expression of *reaper* and *grim* (Fig.3K and K’). In the doomed hemilineage 5A we see robust expression of *grim* (Fig.3I) however, unlike hemilineages 4A, 10A and 12A, we do not see expression of *reaper* (Fig.3I) suggesting that *reaper* is not expressed within this doomed hemilineage population. In lineages where PCD targets hemilineage-B specifically, we find all of the ‘doomed’ hemilineages screened express both *reaper* and *grim* when rescued (Fig.3L-N’). Hemilineage 0B (Fig.3L and L’), hemilineage 14B (Fig.3M and M’) and hemilineage 20/22B (Fig.3N and N’) all display expression of both Reaper and Grim when PCD is rescued in these populations. In summary this tells us that A and B hemilineages as a whole do not exclusively use a single RHG gene to regulate PCD. Most A and B hemilineages, with the apparent exception of 5A, express and utilise both Grim and Reaper.

Interestingly, our T2A-GAL4 reporter alleles also reveal low-level *reaper* and *grim* expression in some of the non-doomed, surviving, sister hemilineage populations. This was evident in lineages 4 (Fig.3H and H’) and 12 (Fig.3K and K’) where the surviving hemilineage B neurons appear to express low levels of both *reaper* and *grim*. In the normally surviving hemilineage 14A (Fig.3M’) low-level *reaper* expression was also observed however this is not accompanied by clear ‘low-level’ *grim* expression (Fig.3M).

In summary, the T2A-GAL4 knock-in alleles of *reaper*, *hid* and *grim* reveal unique spatiotemporal patterns of expression in the late larval CNS. When MARCM lineage analysis is used with the *reaper* and *grim* knock-in alleles in combination with a *dronc^null^* mutation we can observe detailed hemilineage-level patterns of RHG gene expression. These data validate our smFISH data and the T2A cleavage event tells us that they are not only transcribed within these doomed populations but also translated. They show interesting differences in expression and reveal low level expression of *reaper* and *grim* within developing non-doomed thoracic lineages.

### Testing the requirement of Grim and Reaper for hemilineage-specific PCD?

As the expression data show that *reaper* and *grim* are transcribed and translated in doomed neurons soon after they are born. To determine the requirement for both genes in hemilineage-specific cell death we ‘lineage mapped’ postembryonic neurons within the thoracic VNC of L3 larvae with null mutant combinations of *reaper* and *grim* and the deficiency *H99.* To visualise only postembryonic thoracic populations we exploited *worniu-GAL4>UAS-CD8::GFP* (Fig.4A) which labels all neuroblasts and clusters of newly born postembryonic neurons and their stereotyped primary neurite bundles (Fig.4B-D).

**Figure 4:**
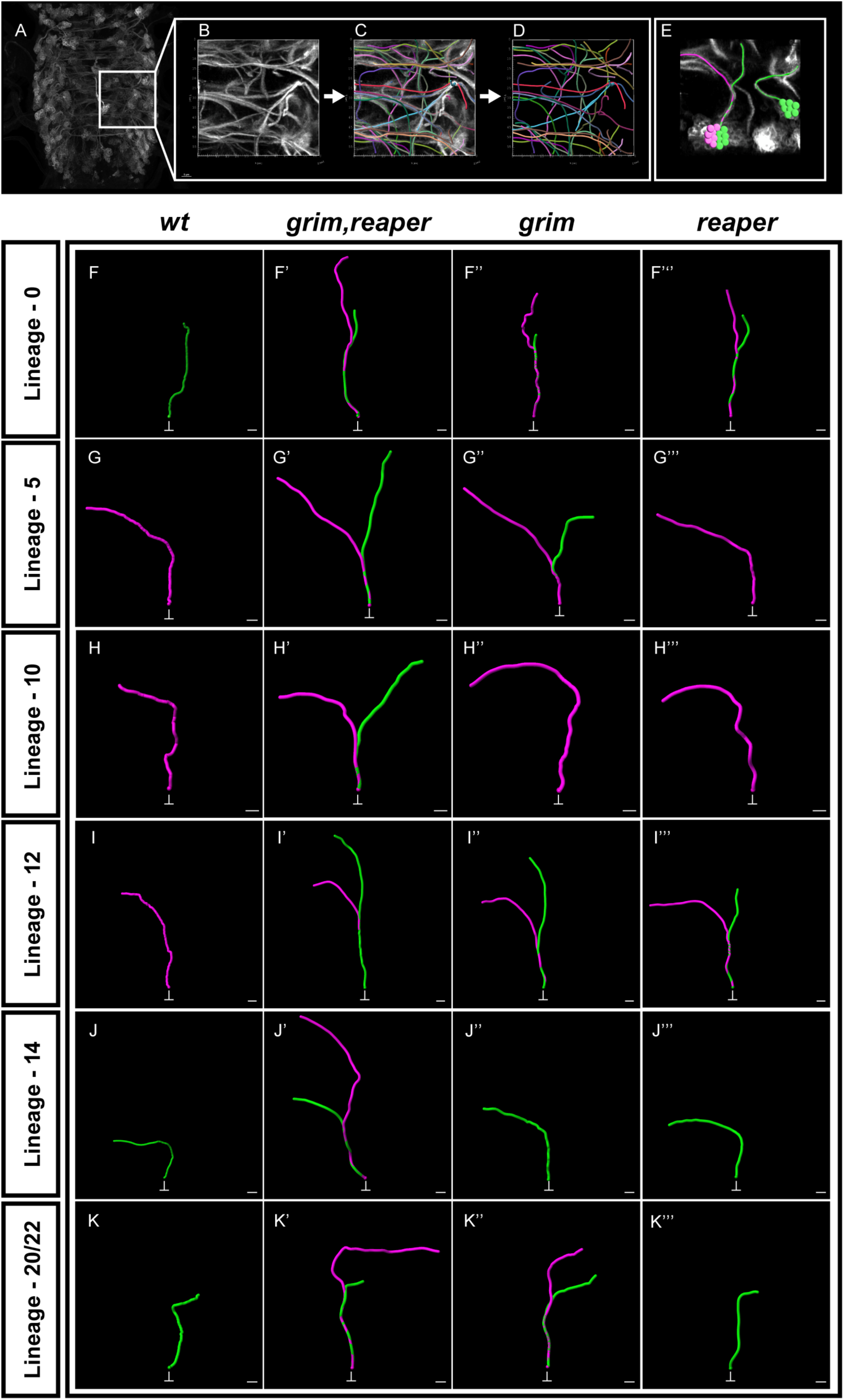
(**A-D**) Workflow of lineage tracing and anatomical model construction. Volumes of thoracic VNC labelled via *Worniu-GAL4* (**A, B**) were traced using the integrated filament module in Imaris. (**C, D**) Traced primary neurites for each lineage were then used to generate an anatomical. Model of the projection pattern of that lineage. Each lineage displayed in **C** and **D** represents a specific lineage. (**E**) anatomical models can trace from lineage origin to termination point and be aligned with know hemilineage anatomy to generate models with hemilineage resolution. (**F – K’’’**) comparison of hemilineage resolution anatomical models of PCD featuring postembryonic lineages from *wt* (n=4), *grim,reaper* (n=7), *grim* (n=8), *reaper* (n=10) individuals at wL3. Lineage 0 (**F-F’’’**). Lineage 5 (**G-G’’’**). Lineage 10 (**H=H’’’**). Lineage 12 (**I-I’’’**). Lineage 14 (**J-J’’’**). Lineage 20/22 (**K-K’’’**). Scale bar, 5μm.

By digitally reconstructing the thoracic lineages we can uniquely identify their respective hemilineages in whole CNSs and determine when cell death is blocked in different backgrounds (Fig.4B-E, Supp Movie.1). For example, in wildtype animals *worniu-GAL4* shows Lineage 6 (where there is no PCD) has two neurite bundles emerging from the cell body cluster with the hemilineage A bundle extending dorsally, crossing the midline in the pD commissure. Hemilineage Bs bundle similarly crosses the midline in the pI commissure and then projects anteriorly (Fig.4E). With Lineage 20/22, a single bundle lineage, where there is PCD, we see a short, dorsal/lateral projecting hemilineage A neurite bundle (Fig.4E). To assess the requirement of *reaper* and *grim* we focused on two sets of lineages, ones where hemilineage-A neurons undergo PCD (5, 10 and 12 in T3) and others where hemilineage-B neurons die (0, 14 and 20/22) (Fig.4F-K).

In ‘single-bundle’ lineages we see the loss of both *reaper* and *grim* together results in a complete blockade of hemilineage-specific PCD in the doomed population. In such cases the normal ‘wild-type’ the neurite bundle are obvious and accompanied by the rescued ectopic bundle (Fig.4F-K). The anatomy and projection patterns of each ectopic bundle matches the previously described anatomy of doomed hemilineage populations demonstrating targeted rescue of the hemilineage population normally fated for removal by PCD (Truman *et al*., 2010).

Specifically for lineage 0 the removal of both *reaper* and *grim* results in an ectopic dorsally projecting neurite bundle extending beyond the termination domain of the normally surviving hemilineage A neurons (Fig.4F-F’). In lineage 10, where hemilineage A dies, we see the loss of *reaper* and *grim* results in a novel laterally projecting bundle of neurites (Fig.4H-H’) in the rescued A hemilineage population. Similarly, in lineage 14, the loss of *reaper* and *grim* blocks death and results in a dorsally projecting lineage bundle (Fig.4J-J’) relating to the rescued hemilineage A population.

Reaper and Grim proteins are similar in structure, both containing the conserved IBM and GH3 domains which characterise the RHG genes (Zhou, 2005). Although there may be specific differences with their binding to IAP proteins, previous studies suggest that both genes are functionally equivalent and capable of killing cells when expressed ectopically (White *et al*., 1996, Chen *et al*., 1996). These observations suggest that any differential requirement may result from differential expression or expression levels for each gene. To determine if there is a division of labour between these two genes when executing hemilineage-specific cell death we looked at how single mutants impact PCD during lineage development.

There are clear examples of lineages where the removal of either *grim* or the removal of *reaper* appears to rescue the doomed hemilineage. Death in Lineage 0 (Fig.4F-F’’’) and the segment-specific death in Lineage 12 (Fig.4I-I’’’) fall into this category, but in 0 this rescue is only partial, i.e. only a small number of doomed neurons are seen which do not extend a full dorsally projecting neurite bundle upon rescue (Fig.4F-F’’’ and Supp Movie. 2). We also find examples of lineages where the doomed hemilineage requires the expression of a single RHG gene for effective regulation of PCD. In lineage 5 we see that removal of *grim* alone results in the rescue of an ectopic hemilineage population, the doomed hemilineage 5A as is seen when both *reaper* and *grim* are removed together in combination (Fig.4G-G’’’). Lineage 20/22 shows a predominant requirement for *grim* expression with removal of *grim* alone manifesting rescue of the doomed hemilineage B (Fig.4K-K’’’).

These data support the idea that *reaper* and *grim* drive hemilineage-specific PCD within the thoracic lineages and that there is a differential requirement in specific hemilineage populations where expression of both or just one are required for mediating sufficient pro-apoptotic drive.

### Temporal patterning of a lineage by RHG-mediated hemilineage programmed cell death generates segment and sex-specific circuit differences

Differences in numbers of specific network components along the neuraxis appears critical to segmental function and the division of labor. Within the thoracic system of *Drosophila* larva ‘monotypic’, single bundle postembryonic lineages, represents the overt morophogical trace of hemilineage-specific PCD sculpting lineages. The removal of an entire hemilineage can be differentially deployed to generate segment-specific differences in circuitry as with, for example, Lineage 1 and hemilineage 12A in T3 (Fig3 K,K’) (Truman *et al*., 2010). One aspect of patterning by deletion we wanted to explore was whether hemilineage-based PCD is differentially deployed in a temporal manner to generate segmental differences in network components in the VNC. Temporal patterning of cell identity within neuroblast lineages appears to be a fundamental feature of cell fate specification in insects (Taghert *et al.,* 1984; Isshiki *et al.,* 2001). To look at temporal aspects of death within hemilineages we focused on later stages of neurogenesis, from larval through to pupal stages and took advantage of the driver R24B02-GAL4 which has been shown to robustly label the hemilineage 12A population during development (Mellert *et al.,* 2016) (Fig.5A). We also chose to investigate patterns of death within this population in both males and females to assess whether changes in PCD deployment occurs in a sex-specific manner during neurogenesis.

**Figure 5:**
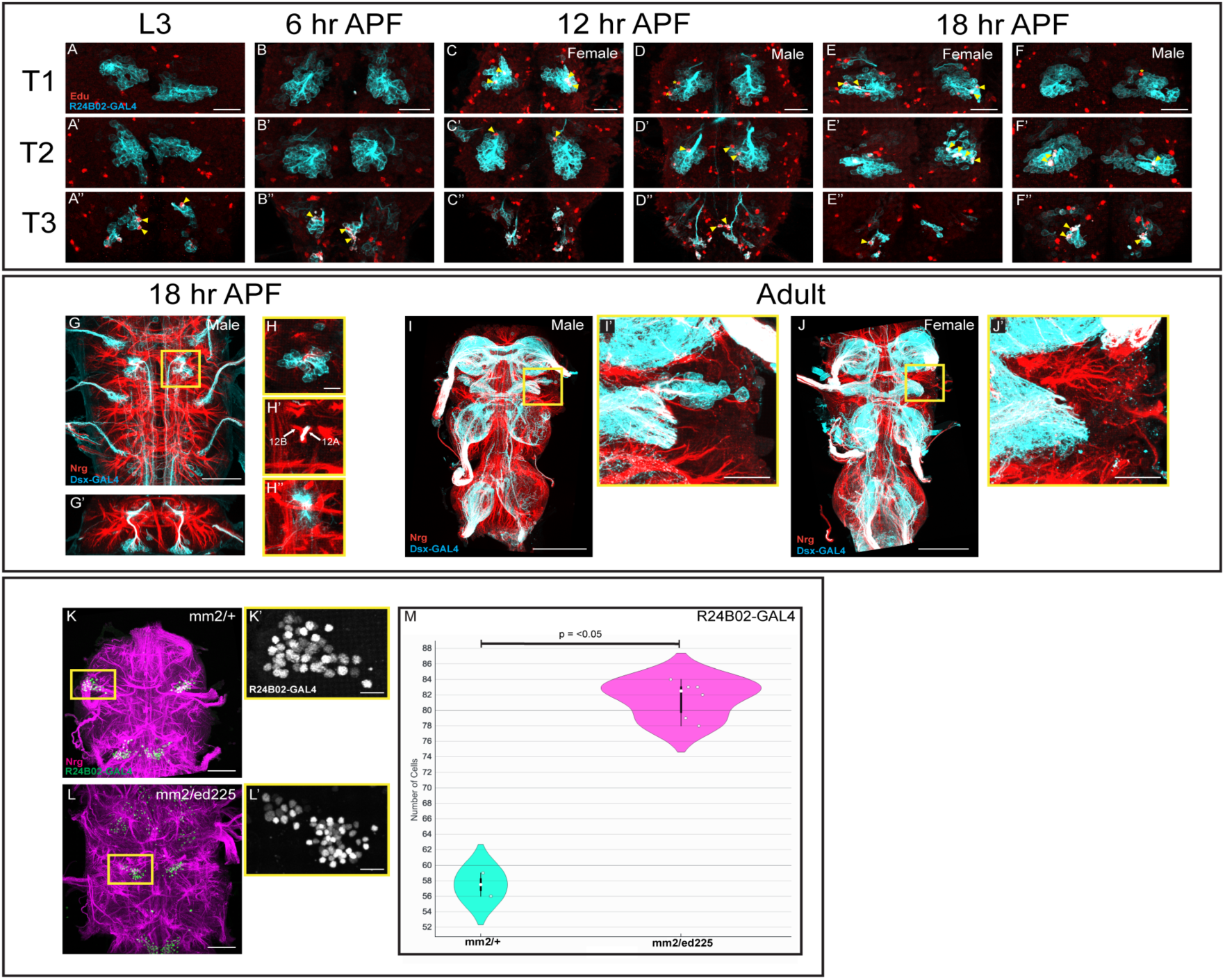
(**A-F’’**) Timeline of active DCP-1 immunoreactivity in hemilineage 12A neurons labelled using R24B02-GAL4 located in T1 (**A-F**), T2 (**B-F’**) and T3 (**A’’-F’’**) at L3 (**A-A’’**), 6 hours APF (**B-B’’**), 12 hours APF females (**C-C’’**), 12 hours APF males (**D-D’’**), 18 hours APF females (**E-E’’**) and 18 hours APF males (**F-F’’**). DCP-1 positive cells located within hemilineage-12A show with solid yellow arrows. DCP-1 positive cells adjacent to, but outside of, hemilineage-12A shown by yellow asterisk. Scale bar, 20μm. (**G-H’’**) Doublesex-GAL4 expression in the male VNC at 18 hours APF. (**G-G’**) Doublesex-GAL4 expression labels a discrete population of dorsally projecting neurones in T1 of the male VNC (yellow ROI). Scale bar, 50mm. (**H-H’’**) Projection pattern of male specific Doublesex-GAL4 positive neurons within the neuroglia scaffold from ventral (**H**) to dorsal (**H’’**) showing projection pattern of sister hemilineages 12A and 12B. Scale bar,10mm. (**I-J’**) Expression of Doublesex-GAL4 in the adult VNC of male and female animals. (**I-I’**) Expression of Doublesex-GAL4 labels a discrete population of Doublesex positive neurons in T1 of the adult male VNC. Scale bar **I,** 100mm. Scale bar **I’,** 20mm. (**J-J’**) Expression of Doublesex-GAL4 showing absence of Doublesex positive neurons in T1 of the adult female VNC. Scale bar **J,** 100mm. Scale bar **J’,** 20mm. (**K-L’**) Labelling of hemilineage 12A in the in the VNC of wt (**K-K’,** mm2/+) and Grim,Reaper mutant (**L-L’,** mm2/ed225) females at approximately 72 hours APF. Scale bar, 50μm. (**K’** and **L’**) T1 associated hemilineage population shown by yellow ROI in **K** and **L** shown in greyscale. Scale bar, 10μm. (**M**) Number of hemilineage-12A neurons in T1 of female VNC at approximately 72 hours APF in wt (mm2/+, n=2) and Grim,Reaper mutant (mm2/ed225, n=6) animals.

During the third larval instar we see PCD within Lineage 12A in the third thoracic neuromere (T3). When death is revealed in labelled neurons using an antibody against cleaved effector caspase DCP-1, the nascent primary neurites extend a short distance from the cell body before dying and fragmenting (Fig.5 A’’). As previously suggested (Marin *et al*., 2012) we do not see death within other 12A populations in the first (T1) and second thoracic (T2) neuromeres (Fig.5 A-A’’). At 6h APF (after puparium formation) the pattern of death within all three segments is the same as seen in the third instar L3, with only 12A neurons in T3 undergoing PCD (Fig.5 B-B’’). At 12h APF we see change, T3 hemilineage 12A neurons continue to be removed (Fig 5 C’’ and D’’) but DCP-1+ve signal is now also evident in the 12A population within the first and second thoracic segment (Fig 5 C’ and D’). Interestingly, at this stage of pupariation we begin to see DCP-1+ve cells only in T1 of females, and not males (Fig 5C and D). This sex-specific pattern of death continues at 18h APF in T1 of females (Fig 5E and F) alongside the non-sex-specific T2 and T3 hemilineage 12A death (Fig.5 E’-F’’’’). The pattern continues >24h APF, until the neuroblasts decommission and neurogenesis ends (data not shown).

As this late induction of T1 death is sexually dimorphic we looked to see if the transcription factor Doublesex, a key regulator of somatic sexual differentiation, is expressed within 12A. Using a GAL4 knock-in into the *doublesex* locus (*dsx-GAL4*) (Manoli *et al*., 2005) to drive a membrane reporter allows the thoracic neuroanatomy in males to be clearly resolved at 18h APF. In males *dsx-GAL4* labels a cluster of dorsally projecting postembryonic neurons in T1 (Fig.5G) that send neurites through a medial bundle in the Neuroglian scaffold (Fig.5G’) bifurcating at an intermediate level and projecting dorsally through the Neuroglian 12A bundle (Fig.5H-H’’). This population of neurons continues to express *dsx* throughout the pupal-adult transition and can be found in eclosed male adult VNC with its bundle of primary neurites passing through the 12A Neuroglian bundle (described in Shepherd et al., 2016) (Fig.5I,I’’). Previous work has referred to this *dsx* +ve population as the TN1 cluster (Lee *et al*.,2002, Rideout *et al*., 2010) There is no *dsx* expression in cells in the same location as the TN1 cluster in females (Fig.5J,J’). Thus late-born members of 12A (*dsx* +ve in adult males) appear to be removed by hemilineage-based PCD in females from 18h APF onwards. To establish if this temporal patterning of 12A death is dependent on Reaper and Grim we removed both genes with the deficiencies MM2 and ED225. To reveal Lineage 12A in both males and females we used R24B02-GAL4 driving the nuclear localising RFP reporter *UAS-RedStinger* (Barolo et al., 2004) (Fig 5K-L’). When counting the number of postembryonic hemilineage 12A neurons generated in T1 at 72 hours APF (∼48 h after neurogenesis has terminated) we see a significant increase in T1 of *grim*, *reaper* mutant females compared to wt (mm2/+) female animals (Fig 5M). Removal of the pro-apoptotic RHG genes *reaper* and *grim* appears to rescue normally doomed hemilineage 12A cells fated for PCD in T1 of the female VNC.

In summary, these data support the idea that Reaper/Grim-mediated hemilineage cell death is critical for sculpting sex-specific circuits during development and that such hemilineage-based ‘temporal patterning’ during neurogenesis maybe an important and widespread means of sculpting segment-specific network motifs.

## Discussion

During nervous system development a panoply of cell fate specification mechanisms generates a large diversity of cell types that assemble into complex networks (ref). One cellular process deployed to control fate in developing nervous systems is programmed cell death by apoptosis (Buss *et al.,* 2006). The majority of studies in insects have focused on the death of fully differentiated neurons either at the onset of metamorphosis or following adult eclosion (Truman *et al.,* 1992). In previous work we revealed that the most common type of cell death within the developing insect nervous system is hemilineage-based PCD that occurs during neurogenesis. Hemilineage-specific PCD is precisely patterned and removes immature neurons, soon after they are born, ensuring that populations of specific neuronal subtypes are generated in the right numbers, in the right places. During neuronal development cell fate specification mechanisms result in type-specific combinations of transcription factors, called ‘terminal selectors’, that initiate and maintain terminal identity programs by direct regulation of subtype-specific ‘effector genes’(Hobert, 2016). Until now the pro-apoptotic genes that act as the terminal ‘effector genes’ directly driving hemilineage-specific death have not been determined. Here we show how RHG gene expression maps onto developing lineages of *Drosophila* and that their expression controls PCD within hemilineage populations.

### Grim and Reaper are the ‘effector genes’ for hemilineage-specific PCD fate

Clonally removing *reaper*, *hid* and *grim*, together, completely blocks hemilineage-specific PCD in postembryonic neurons. These *H99* deficient lineages, typically have ectopic neurites emerging from the ‘rescued’ hemilineage, along with the neurites from the ‘wildtype’ non-doomed sister hemilineage. These lineages with undead neurons look like the clones generated with *Dronc* caspase null mutants (Truman *et al.,* 2010). Our RHG smFISH data show *reaper* and *grim* transcripts in discrete clusters in dividing neuroblasts and new-born postembryonic throughout the thoracic VNC but find no evidence for *hid* or *sickle* transcripts in these same cells. <u>Our lack of observation of *hid* and *sickle* expression suggests that these RHG genes are not utilised during the specification of hemilineage PCD in the VNC. This suggests that the transcriptional cues specifying death upstream of the RHG genes are specifically regulating only the expression of *grim* and *reaper* in the VNC hemilineages.</u>

In dronc null mutants we see large numbers of undead cells that have robust *reaper* and *grim* expression, suggesting that hemilineage-specific PCD is the result of their transcription in doomed hemilineage populations. This is supported by a number of functional studies that show ectopic induction of *reaper* and *grim* transcription is capable of driving cells to apoptosis (Grether *et al*., 1995, Chen *et al*., 1996, Nordstrom *et al.,* 1996, White *et al.,* 1996). The sustained expression of *reaper* and *grim* transcripts within these undead*, dronc^-^* null, neurons suggests that the RHG genes are equivalent to cell-type specific ‘effector/terminal identity’ genes (Hobert, 2016). Which, in this case, regulate the adoption and maintenance of the fate of death within VNC lineage neurogenesis.

### Revealing hemilineage-specific patterns of Reaper and Grim expression

The MARCM clonal analysis with our T2A-GAL4 knock-in alleles, in combination with the *dronc* null allele, reveals hemilineage-specific expression of both *reaper* and *grim* in these doomed populations. As well as validating the smFISH observations the T2A-GAL4 clone data provides two key observations. First, self-cleaving nature of the T2A fusion reagents also tells us that the RHG transcripts are being actively translated within the doomed hemilineage populations at this time. More importantly it also enabled us to visualise neurite anatomy of individual populations and gain hemilineage-level resolution of RHG gene expression.

One idea we addressed was whether there is mutually exclusive Notch-dependent RHG expression within doomed lineages as Bertet and collegues show that in the developing optic lobe *hid* and *reaper* regulate the apoptotic fate of specific hemilineage progeny (Bertet et al., 2014). There, Notch-ON and OFF death is underpinned by temporal transcription factor expression, with Notch-ON death in early progeny being Reaper dependent and Notch-OFF death in late progeny requiring Hid. In the VNC populations described here we found that *reaper* and *grim* appear to be expressed together, with no clear indication of hemilineage-A (Notch-ON) and hemilineage-B (Notch-OFF) cells differentially expressing either *reaper* or *grim* alone. Additionally, we never see *hid* expression in the doomed postembryonic VNC neurons, even in the *dronc^-^* null background but we do observe *hid* expression within the optic lobes (Fig.3.B) with the *hid-T2A-GAL4* allele. These data, and that of others, suggest that spatiotemporal patterning of *reaper*, *hid* and *grim* expression during development is diverse, with the locus integrating different regulatory inputs to orchestrate death with in different cellular contexts, in different tissues at different times.

Our T2A reporter tools also revealed low-level expression of *reaper* and *grim* in many cell types in the CNS (Fig 3. B-D) even within ‘non-doomed’ hemilineages. This low-level *reaper/grim* expression in non-doomed hemilineages suggests that some hemilineages that do not normaly die express *reaper* and *grim* but at levels below the ‘apoptotic threshold’ (Florentin and Arama, 2012). Alternatively, this expression may play some other role in lineage development, as we know RHG proteins enable physiologically relevant non-apoptotic caspase function in many cellular contexts (Nakajima and Kuranaga, 2017). A more mundane explanation could be that these hemilineages are ones we have assumed to be surviving but are switching into hemilineage-based PCD at this stage of development, although we think this unlikely.

### The differential requirement of Reaper and Grim within hemilineages

The complexity of expression we see within thoracic lineages for *reaper* and *grim* suggests a divergence in their function. To address this, we tested their requirement by removing *reaper* and *grim* individually and in combination. The loss of both genes appears to block all hemilineage-specific PCD within the postembryonic VNC lineages supporting the hypothesis that they are the primary pro-apoptotic effector genes driving this mode of death. Interestingly, when removed individually we see complex differential requirements, suggesting the presence of nuanced, lineage-specific regulatory mechanisms.

Our expression and functional data (Fig. 4) along with previous work on these genes suggests that this locus is under complex spatiotemporal transcriptional control and that the landscape of regulatory modules surrounding *reaper* and *grim* orchestrate distinct hemilineage-specific patterns of transcription. Previous studies on the transcriptional control of these pro-apoptotic genes have shown them to play a key role in the controlling of neuroblast death during postembryonic development (Tan *et al.,* 2011, Bello *et al*., 2003, Khandelwal *et al*., 2017,). A 20-kb regulatory region between *reaper* and *grim* called the Neuroblast Regulatory Region is responsible for the PCD of abdominal neuroblasts (Tan *et al.,* 2011). Elegant studies dissecting the transcriptional control of the NBBR reveals discrete *cis*-regulatory modules each containing multiple transcription factor (TF) binding sites (Tan *et al*., 2011, Arya *et al*., 2015). Could a similarly complex *cis*-regulatory control exists for hemilineage-based PCD as each thoracic hemilineages expresses distinct combinations of putative ‘terminal selector’ transcription factors (Lacin *et* al., 2014, Lacin and Truman, 2016; Allen *et al*.,2020).

By identifying the two RHG genes regulating this precisely patterned PCD we have clear candidates to focus on future investigations of how the surrounding genomic regulatory landscape directs their transcription.

### Temporal patterning of a hemilineages generates segment and sex-specific circuit differences

Differences in the numbers of specific neuronal subtypes along the neuraxis is a critical feature of network organization and function. To explore this, we analyzed whether there is evidence of segment specific temporal deployment of hemilineage PCD in thoracic neurogenesis. Temporal patterning presents a mechanism to generate differences in neuronal numbers by varying when PCD begins to eliminate neurons within segmental lineage homologs. Such rank order-based cell fate specification mechanisms are a fundamental feature of insects CNS development (Pollington et al.,2023). To look at temporal aspects of PCD we focused on a single hemilineage throughout neurogenesis and find that in late stages sex-specific patterns of greater PCD are seen in the T1 hemilineage 12A population in females. The eliminated neurons appear to be the male-specific Doublesex-positive neurons known to be critical for male courtship song in *Drosophila* (Rideout *et al.,* 2010; Shirangi *et al.,* 2016). The changes in the patterns of cell death between segments and sex show how malleable the system can be. It appears that each hemilineage exists as an individual development unit that can be manipulated in isolation to effect changes in circuit development and function. We anticipate that there may be other lineages with variations in population size throughout the neuraxis that arise from differential deployment of hemilineage PCD during neurogenesis. We know that the Hox gene *Ubx* controls hemilineage specific PCD in T3 to generate the monotypic single bundle lineage 12A found in that neuromere (Marin *et al*., 2012) and that Hox inputs into the transcription regulation of *reaper* and *grim* are evident. It is possible that different Hox genes which pattern the A-P axis are important inputs regulating expression of *reaper* and *grim* within specific temporal hemilineage cohorts along this axis of the VNC.

In the context of sexual differences within the nervous system it seems that RHGs play a significant role (Kimura *et al*., 2005). Ghosh and colleagues show AbdB and Dsx^F^ undergo cooperative binding at an enhancer in the *reaper* locus in females but not in males. Neuroblasts undergo apoptosis in females whereas the equivalent counterparts in males proliferate, giving rise to serotonergic neurons that are important for adult mating behaviour (Ghosh *et al*.,2019). We predict that differences in hemilineage-specific PCD, mediated by distinct terminal selector gene combinations result in the differential transcription of RHGs that play a role in regulating the numbers of different neuron types and network motifs in males and females.

### Differential regulation of RHG expression and the evolution of novel network motifs

A key take-home from our work is the how this modular control of hemilineage cell death by RHG genes can play out in an evolutionary context. Insects are arguably the most diverse and speciose group of animals in the biosphere. They show a high degree of morphological variation and diverse behavioural repertoires but remarkably the developmental anatomy of the CNS remains highly conserved with the segmental array of 61 neuroblasts being strikingly stable across the order (Truman and Ball, 1998, Biffar and Stollewerk, 2014). We propose that that hemilineages may be key ‘evolvable modules’ of change (Pop *et al*.,2020).

The differential requirement of both genes and the sub-threshold patterns of RHG expression we report here provides a framework that points to how such changes could occur. Within a hemilineage there may be a set level of RHG expression/apoptotic drive and that dialling this up or down within that population may shift populations incrementally towards the ‘apoptotic threshold’ (Florentin and Arama, 2012). In this way changes in the number of network components for building the adult CNS can appear. If novel hemilineage-specific PCD patterns emerge and are selected upon they could facilitate the emergence of novel network motifs in species occupying different ecological niches. Similar changes can occur within the sensory system where altered patterns of PCD may result in novel sensory neuron types emerging (Prieto-Godino *et al.,* 2020). In support of this, we have observed that in dipterans that have evolved secondary wing-loss, i.e. are now flightless, there is strong neuro-anatomical evidence to suggest that populations known to be involved in flight may be targeted for removal by hemilineage PCD (Pop et al., 2020, Sproston and Williams, unpublished). We hope that through further comparative studies we will be able to explore this idea and address whether changes in the regulatory inputs on to the RHG locus form a key node through which evolution of the CNS may take place.

## Supplemental Figures

**Supplemental Figure 1:**
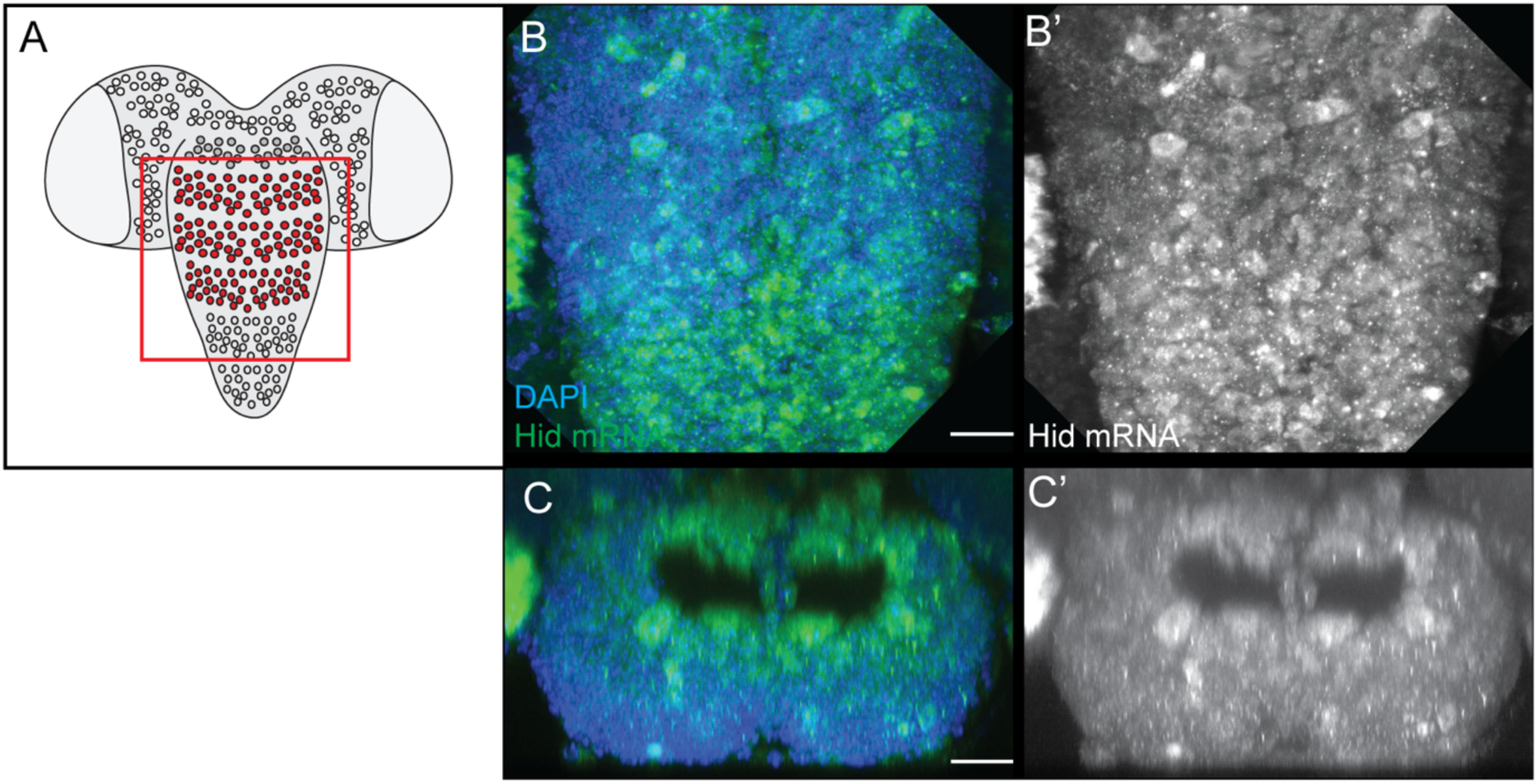
Heatshock induced Hid expression in the VNC at wL3 stage **(B-C’). (B)** Schematic showing overview of late larval CNS. ROI featured in data panels **B-C’** shown in red boxed region. (**B**) maximum intensity projection of ventral region of VNC showing heatshock induced Hid mRNA expression (green) and DAPI (blue). (**B’**) Greyscale image of Hid mRNA expression shown in **B. (C)** Transverse maximum intensity projection showing heatshock induced Hid mRNA expression in the thoracic VNC. Hid mRNA (green), DAPI (Blue). **(C’)** Greyscale image of Hid mRNA expression shown in **C.** Scale bar, 20 μm.

**Supplemental Figure 2:**
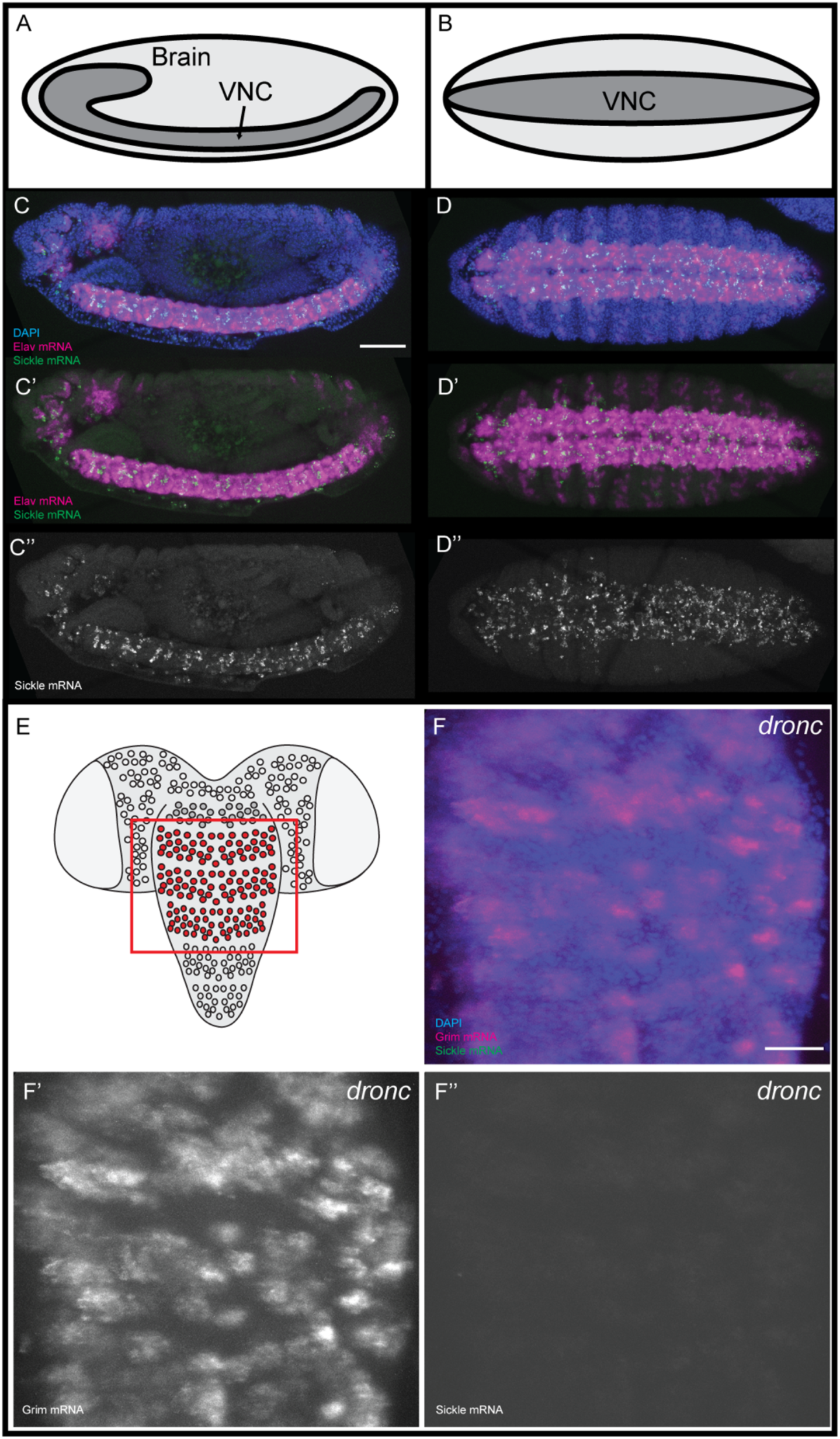
Sickle mRNA expression in the embryonic CNS and doomed hemilineages of the late larval VNC. (**A**) Schematic of lateral view of embryo as shown in **C-C’’** showing rough anatomy and position of CNS including Brain and VNC. Anterior = left, posterior = right. **(B)** Schematic of ventral view of embryo as shown in **D-D’’** showing rough anatomy and position of the VNC. Anterior = left, posterior = right. **(C-C’)** Lateral view of Elav and Sickle mRNA expression in the embryo at ∼stage 15. **(D-D’)** Lateral view of Elav and Sickle mRNA expression in the embryo at ∼stage 15. DAPI signal **(C** and **D)** demarcated boundary of embryo proper. Elav mRNA expression **(C-C’, D-D’)** demarcates boundaries of the CNS/PNS at stage 15. Scale bar, 50 μm. (**E**) Schematic of late larval VNC. Red box shows region of thoracic VNC displayed in **F-F’’. (F-F’’)** Thoracic VNC of *dronc^-^* larval VNC showing expression of Grim mRNA (**F-F’**) and Sickle mRNA (**F-F’’**) in the rescued doomed hemilineages. DAPI (**F**) demarcates boundaries of the thoracic VNC. Scale bar, 20 μM.

**Supplemental Figure 3:**
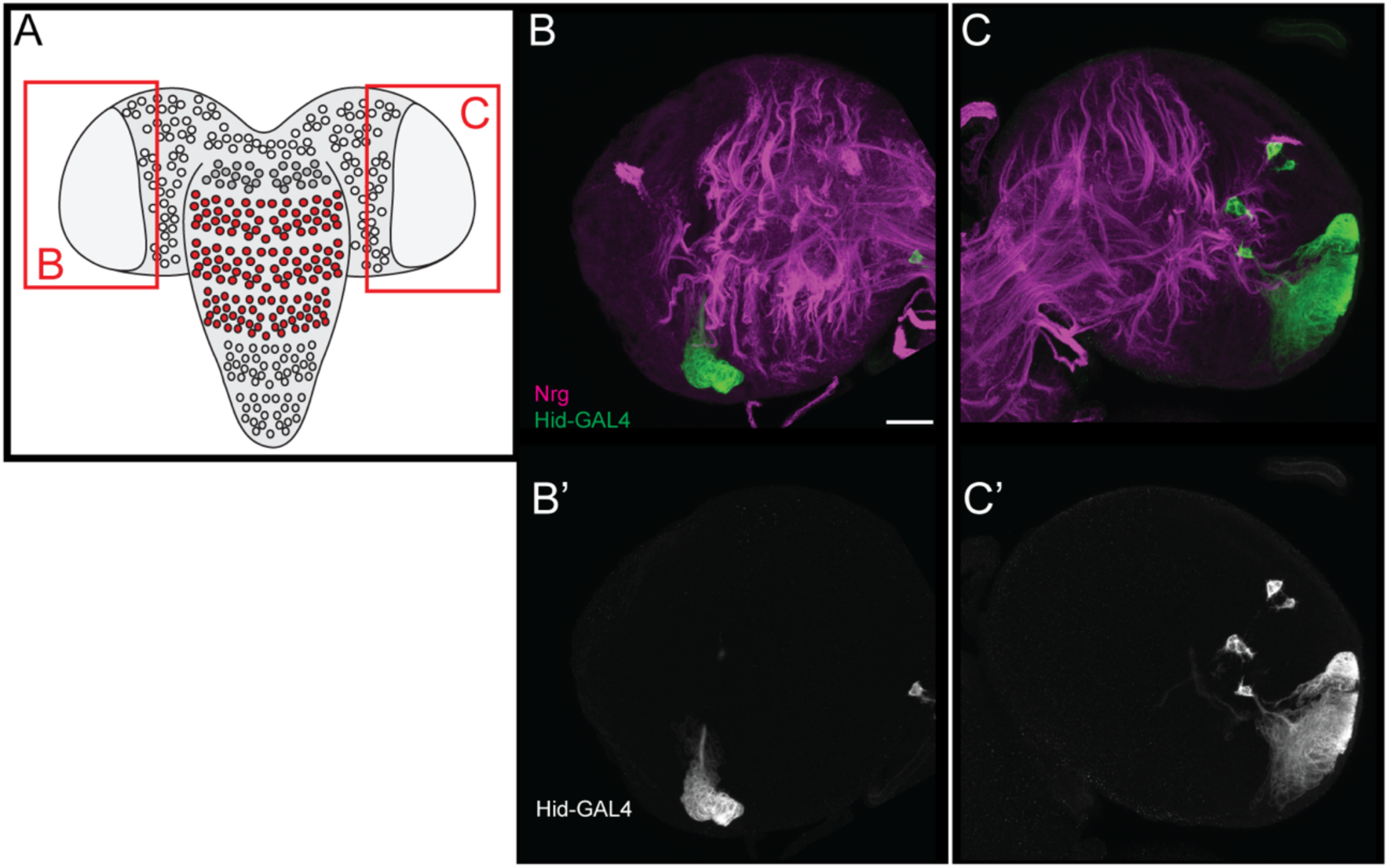
Larval induced Hid-T2A-GAL4 dronc^-^ MARCM clones in the developing optic lobe at wL3. **(A)** Schematic showing anatomy of larval CNS at wL3. Red boxes display optic lobe ROI’s shown in **B-B’** and **C-C’. (B-C’)** Hid-T2A-GAL4 expression MARCM clones induced at newly hatched larval stage and recovered at wL3 showing expression in lineages of the developing adult visual system. Scale bar, 20 μm

**Supplemental Video 1:** Video showing example filament tracing for a single wild-type Lineage (Lineage-0). Left shows VNC volume with traced Lineage-0 model overlaid. Right shows VNC volume without traced Lineage-0 model overlaid. Lineages labelled via expression of UAS-CD8:GFP under the control of Wor-GAL4

**Supplemental Video 2:** Video showing example partial rescue of hemilineage 0B in a GrimA6C/H99 VNC. Green shows location and projection of normally surviving hemilineage 0A. Magenta shows thin projection of partially rescued hemilineage 0B. Lineages labelled via expression of UAS-CD8:GFP under the control of Wor-GAL4

## Materials and Methods

### Image acquisition and processing

All Images were acquired on a Zeiss LSM800 at a magnification of 40x or 63x (where specified) using a 1μm step size. All image analysed and processed using FIJI (https://imagej.net/software/fiji/). For *Drosophila* MARCM clones, lineages were cropped using the freehand tool in FIJI to remove visual contamination of neighbouring lineage clones and displayed as maximum intensity projections.

3d models of larval hemilineages were generated in IMARIS (version: 9.10) using manual tracing with the filament tracer tool. All 3d renders and videos were generated in IMARIS.

All figures were constructed using Adobe Illustrator 2023.

### Data tabulation and plotting

All numerical data was collected and curated in Microsoft excel and plotted using GraphPad Prism^TM^.

### Drosophila husbandry

Fly stocks were raised on standard cornmeal/glucose media. Stocks were maintained at 22 °C on an approximately 12hr:12hr light dark cycle during general stock maintenance and for the collection of virgin and male stock for experimental crosses. All experimental crosses were maintained at 25 °C on a 12hr:12hr light dark cycle in order to allow for the precise timing and staging of larval development.

### Fly stocks used in this study

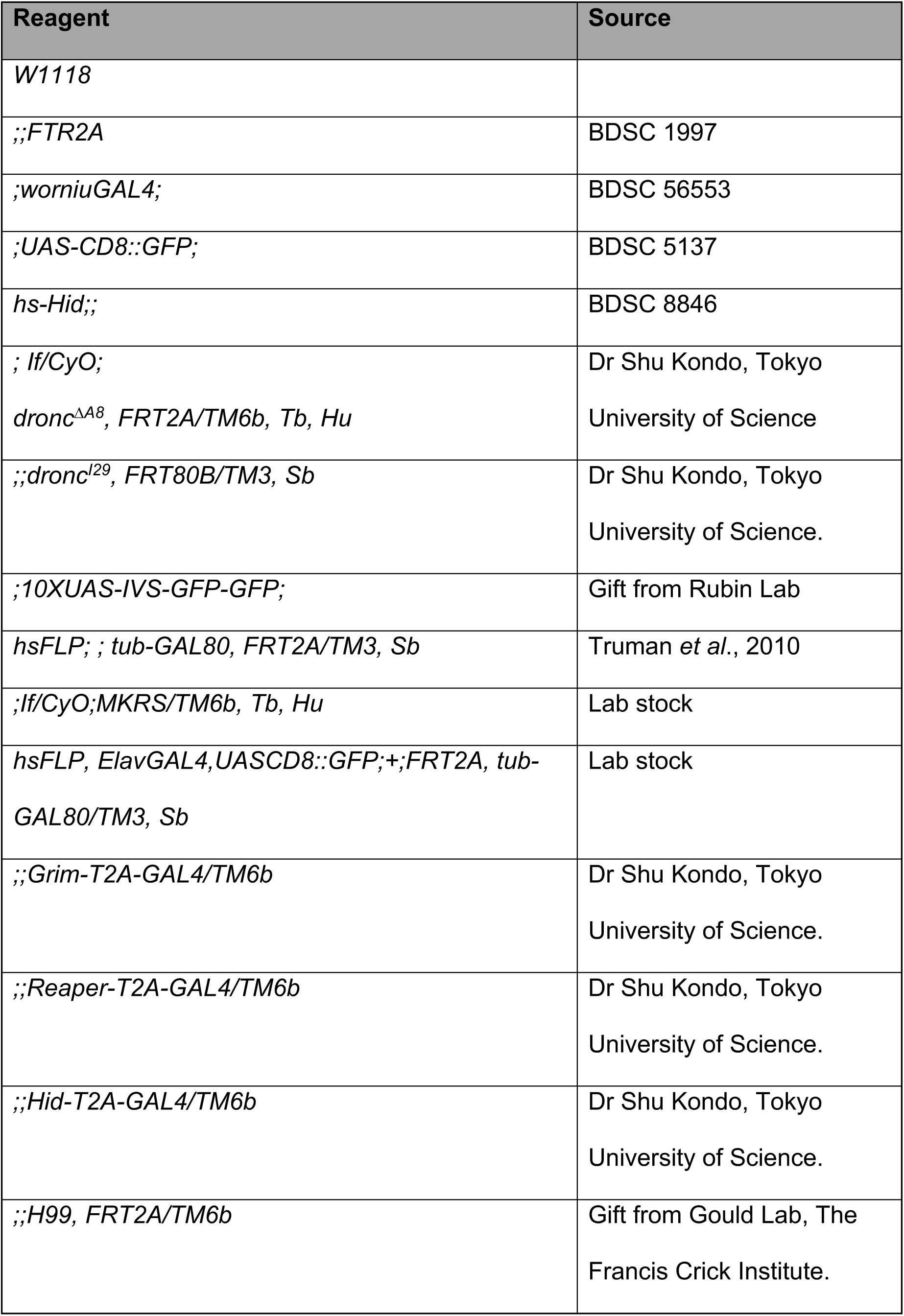
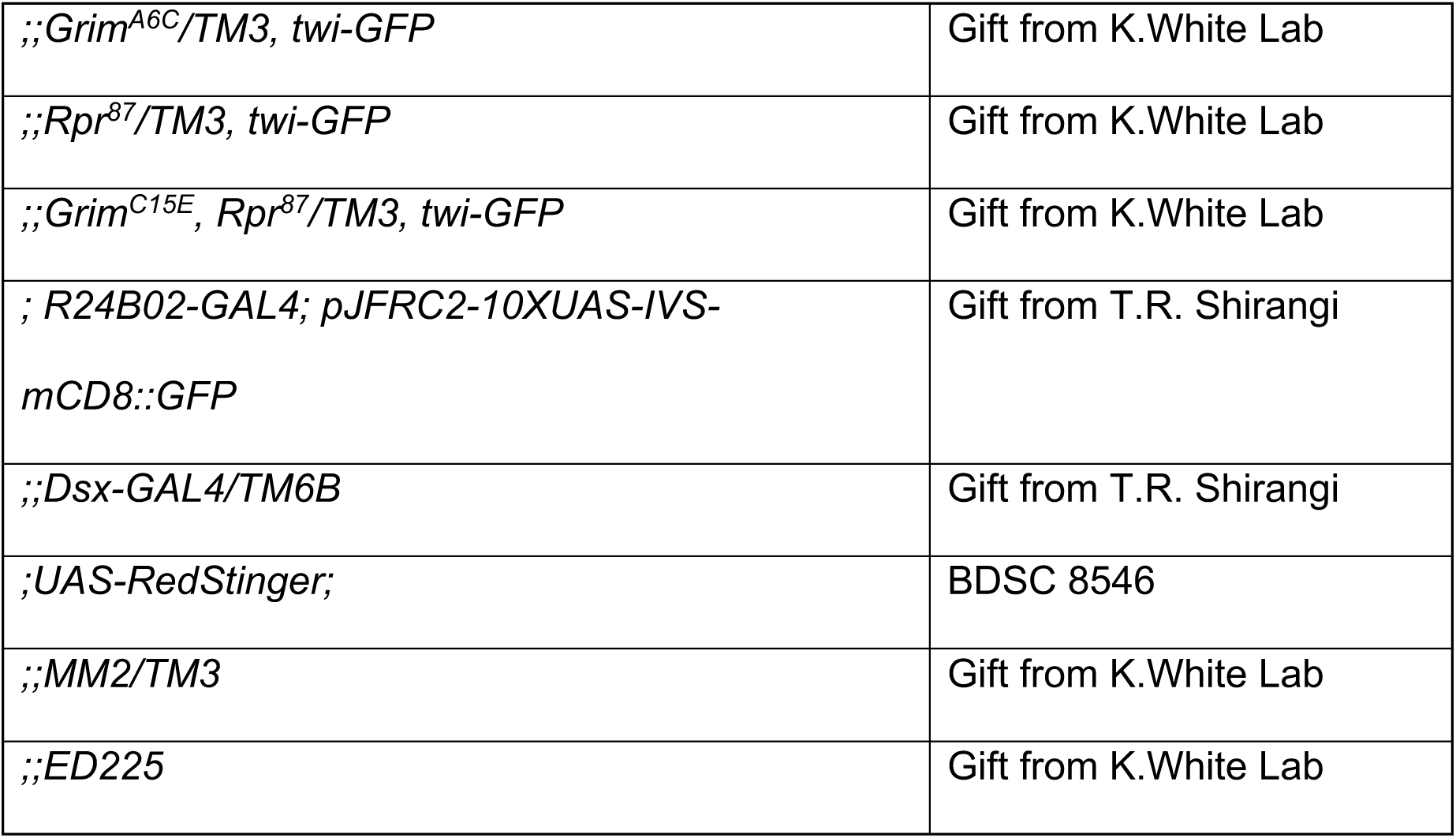

### MARCM clone generation

Mitotic postembryonic lineage clones were generated using the **M**osaic **A**nalysis with a **R**epressible **C**ell **M**arker technique (**MARCM**) (lee and luo., 1999). 0-4hr post hatching 1^st^ instar larvae were heat-shocked at 37 °C in plastic, food containing, vials for 1 hour before being returned to 25 °C in order to develop to 3^rd^ instar. For denser clonal inductions the above process was accompanied by 30 mins at 25 °C before being returned to 37 °C for a further 30 mins.

Dronc MARCM clones were generated by crossing virgin females of the genotype *hsFLP, ElavGAL4, UAS-CD8::GFP;+;FRT2A, tub-GAL80/TM3, Sb* with male flies of the genotype *;;Dronc* ^D^*^A8^,FRT2A/TM6B, Tb, Hu*.

Grim-T2A-GAL4 and Reaper-T2A-GAL4 expressing clone inducible crosses were set up through the mating of virgin females of the genotype *hsFLP;;tub-GAL80, FRT2A/TM3, sb* with males from one of the following MARCM compatible stocks:

- ;*UAS-CD8::GFP/CyO; Grim-T2A-GAL4, FRT2A/TM6B*
- *;UAS-CD8::GFP/CyO; Reaper-T2A-GAL4, FRT2A/TM6B*
- *;UAS-CD8::GFP/CyO; Grim-T2A-GAL4, dronc*^D*A8*^*, FRT2A/TM6*
- *;UAS-CD8::GFP/CyO; Reaper-T2A-GAL4, dronc*^D*A8*^*, FRT2A/TM6B*

### CNS Dissections

Larvae at wandering L3 (wL3) and pupae were collected from food containing vials and their CNS dissected out in ice cold phosphate buffered saline (1xPBS). The CNS was transferred to 1xPBS on ice before fixation. Dissections were carried out within 1 hour. CNS were fixed in ice cold 3.6%PFA in PBS for 1 hour with shaking. After fixation fixative was removed and samples were prepared for immunohistochemical staining.

### Immunohistochemical staining

Fixed CNS’s were first washed for 4 x 15 minutes at room temperature in 0.3% PBST (0.3% Triton X-100 in 1xPBS). After washing samples were blocked for 1 hour at room temperature in blocking solution (5% neutral goat serum in 0.3% PBST). After blocking samples were incubated with primary antibody diluted in blocking solution. Primary antibody incubations were carried out for between 1-3 days at 4 °C. After primary antibody incubation samples were washed thoroughly (minimum 4x throughout the day) in 0.3% PBST. After washing of primary antibody solution, samples were incubated with the appropriate secondary antibodies diluted in blocking solution for a further 1-3 days. After secondary antibody incubation samples were thoroughly washed with 0.3% PBST throughout the day. After a final wash for 30mins in PBS, samples were mounted on poly-L-lysine coated cover slips. Mounted samples were dehydrated through immersion in coverslip jars containing increasing serial concentrations of ethanol. Slides were incubated in 15%, 30%, 70%, 80%, 90% and twice in 100% EtOH for 5 minutes. After EtOH dehydration coverslips were dipped once in Xylene and subsequently incubated twice for 10 minutes in fresh Xylene in coverslip jars. After both incubations a drop of DePeX mounting media was placed over the mounted samples and the coverslip inverted (sample side down) and gently placed onto a clean, labelled, microscope slide. The coverslip was gently pressed/tapped into place before being transferred to a slide box and placed at 2 °C protected from light.

### RHG specific smFISH probe design

Stellaris RNA FISH probes for *D.mel reaper*, *grim*, *hid* and *sickle* were ordered from Biosearch Technologies^TM^(https://www.biosearchtech.com/support/tools/design-software/stellaris-probe-designer). Probe design was carried out using the Stellaris Probe Designer tool and in accordance with the Stellaris probe design guidelines. Probes were designed against the coding sequence (CDS) for each of the RHG genes with a masking level of 5. For Hid probes were designed against only the exon sequences of the longer putative Hid splice variant (Hid-RA). Probe sequences for each RHG were aligned to *D.melanogaster* transcriptome through BLAST in order to check probe specificity. Probes that showed a high complementarity to off-target RNA were omitted. If complete complementation was observed or, if 5 or more probes showing 16 or more nucleotides complementarity to an off-target RNA were observed, those probes were omitted. In order to generate specific probe sets which contain the recommended number of 25+ probes parameters for probe nucleotide length and spacing (nucleotide gap between probes) were adjusted for each RHG gene. If after BLAST analysis a sub 25 probe set was retrieved the above parameters were adjusted and the resulting probes re-analysed through BLAST until an appropriate probe set was designed.

The final parameters for each RHG set were as follows:

- *hid* – Length: 20 Spacing: 2
- *grim* – Length: 22 Spacing: 1
- *reaper* – Length: 18 Spacing: 1
- *sickle* – Length:20 Spacing: 2

### smFISH probe preparation and storage

Probes were resuspended in 95μl fresh TE with 5μl RNase inhibitor (Promega: RNasin Plus RNase inhibitor cat:N2611). 5μl aliquots. Were prepared and immediately transferred to −80 °C for storage.

### smFISH hybridization and mounting of late larval CNS

CNS were dissected at in ice cold 1xPBS (DEPC treated, Filtered) and transferred immediately to 1xPBS on ice. All dissections were carried out within 2 hours. After dissection samples were tubes and fixed for 1 hour at room temperature with nutation in 3.6% PFA (16% PFA diluted in DEPC treated and filtered 1xPBS). Samples were washed in 0.3% PBST (0.3% Triton X-100 in DEPC treated, filtered 1xPBS) for 3x 15 minutes at room temperature with nutation. Samples were then dehydrated in a short ascending ethanol series (15% - 30% - 70% EtOH diluted in DEPC treated and filtered H_2_O). Samples were incubated at room temperature for 10 minutes in 15%, 30% and 70%. Once in 70% EtOH samples were transferred to RNase free 500ul microcentrifuge tubes, topped up with 70% EtOH and incubated overnight at 4 °C with shaking. After overnight incubation EtOH was removed and 100μl of fresh Wash Buffer A (100μl Stellaris® RNA FISH Wash Buffer A with 350μl DEPC treated de-ionised H_2_O, 50μl HiDi Formamide) was added and incubated for 5 minutes at room temperature. After 5 minutes wash buffer was removed and 100μl Hybridization buffer (450μl Stellaris® RNA FISH Hybridization Buffer with 50μl HiDi Formamide) with 250nM of Hybridization probe (0.5μl of each probe per 100μl of Hybridization solution) was added. Samples in Hybridization buffer + probes were incubated for 16 hours at 37 °C with shaking. After hybridization, hybridization buffer + probe solution was removed and samples were washed with wash buffer a 3x for 15 minutes at 37 °C with shaking. After washing samples were incubated with DAPI (1:500 in Wash Buffer A) for 30 minutes at 37 °C with shaking. Wash buffer A (DAPI) was removed and samples were washed with Wash Buffer B (Stellaris® RNA FISH Wash Buffer B) for 5 minutes at room temperature. Samples were washed once in 1xPBS for 5 minutes and mounted ventral side up on poly-L-lysine cover slips and mounted in vectashield. Once mounted samples were stored at −20 °C before imaging.

### smFISH hybridization and mounting of whole embryos

Embryos were then incubated at 25 °C for 12 hours until stage 15-16. Embryos were collected with a paintbrush and dechorionated in 50% bleach for 5-10 minutes. After dechorionation embryos were washed with ddH_2_O and then transferred to a glass scintillation vial containing 5ml 3.2% PFA (37% PFA diluted in DEPC treated and filtered 1xPBS) + 5ml Heptane. Embryos were fixed for 30 minutes before the aqueous PFA layer was removed and discarded. After removal of the aqueous layer 5ml (equal volume) of MeOH was added and the embryo/heptane/MeOH mix was shaken for one minute to remove the vitelline membrane. The de-vitellinised embryos were then pipetted from the bottom of the scintillation vial, transferred to a 500ul microcentrifuge tube and washed thoroughly with fresh MeOH. After MeOH wash, embryos were rinsed twice, for 5 minutes with shaking, in 0.3% PBST (0.3% Triton X-100 in DEPC treated, filtered 1xPBS). After short 5-minute rinses, embryos were washed in 0.3% PBST for 3x 15 minutes at room temperature with nutation. Following PBST wash embryos were dehydrated, hybridised and mounted as detailed above for whole mount CNS smFISH experiments.

### Heat-shock induced *Hid* expression

Using a heat-shock inducible Hid *Drosophila* strain allowing for the driving of Hid expression under the control of the heat-shock promoter Hsp70. The particular stock was a Y chromosome male killing variant of the conditional heat-shock induced Hid strains generated via the remobilisation of the *P{hs-Hid}* transposable element first described by (Grether *et al*., 1995). Third instar larvae were either heat-shocked at 37 °C for 20 minutes. Male larvae were then dissected, fixed and smFISH carried out probing for Hid mRNA expression.

### Antibodies

Antibodies used in this study.

| Species | Antigen | Dilution | Labeles | Source | Citation<sup>*</sup> |
| --- | --- | --- | --- | --- | --- |
| Mouse | anti-Neuroglian | 1:50 | Postembryonic lineage bundles | DSHB<br>BP 104, |  |
| Chicken | anti-GFP | 1:500 | GFP | Abcam<br>ab13970 |  |
| Rabbit | anti-Cleaved *Drosophila* Dcp-1 | 1:100 | Cells undergoing apoptosis (activated effector caspase) | Cell Signalling<br>9578 |  |
| Goat | anti-Chicken 488 conjugated | 1:500 | Secondary Antibody | Invitrogen A11039 |  |
| Donkey | anti-Mouse Cy5 conjugated | 1:500 | Secondary Antibody | Jackson<br>ImmunoReaserch<br>715-006-151 |  |
| Donkey | anti-Rabbit Cy3 conjugated | 1:500 | Secondary Antibody | Jackson<br>ImmunoReaserch<br>711-006-152 |  |

\* Citation supplied if antibody received as gift

### smFISH probes designed as part of this study

List of Grim mRNA specific probe sequences (5’-3’) designed and used in this study:

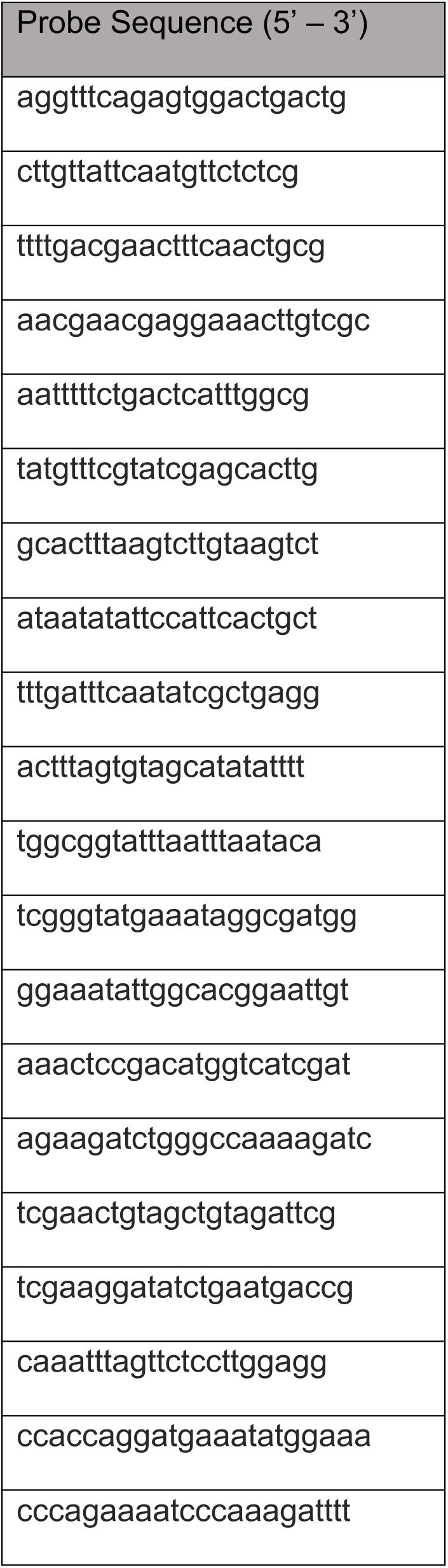
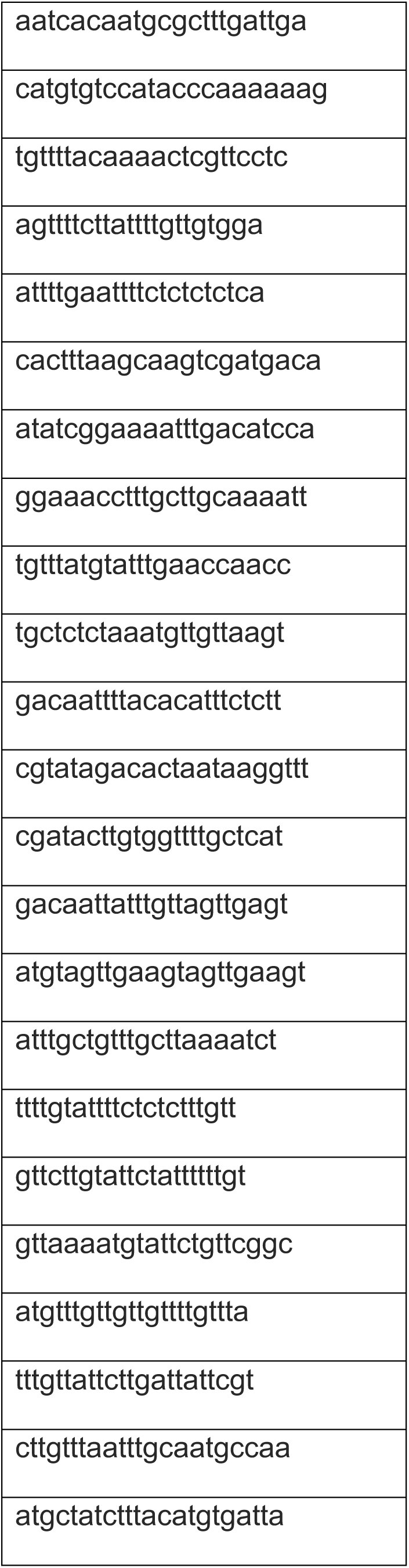

List of Reaper mRNA specific probe sequences (5’-3’) designed and used in this study:

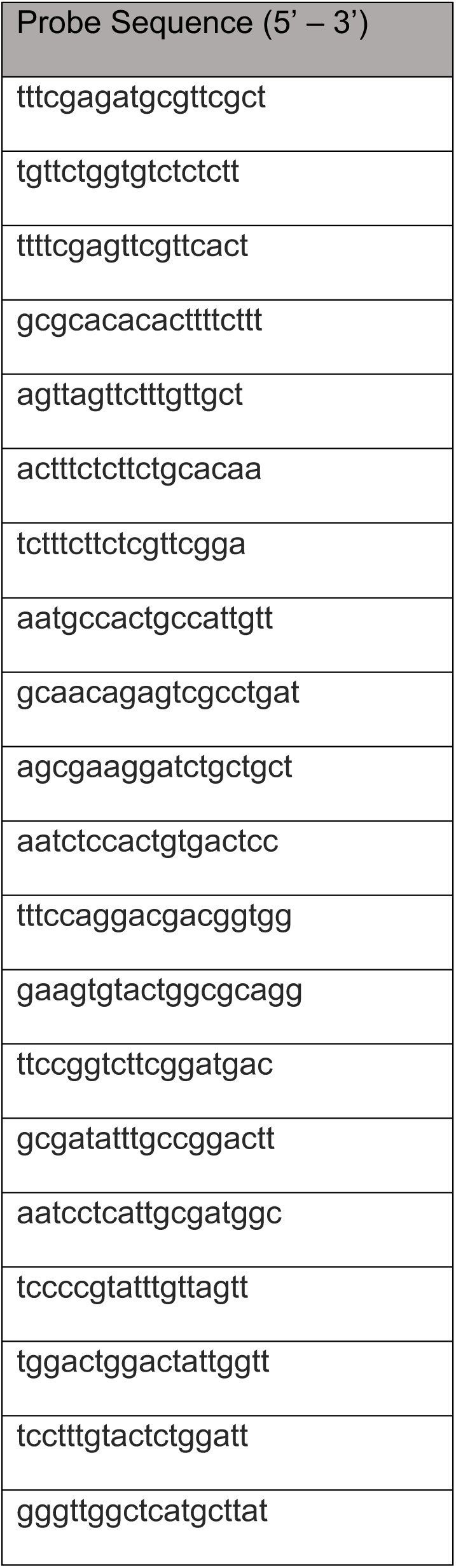
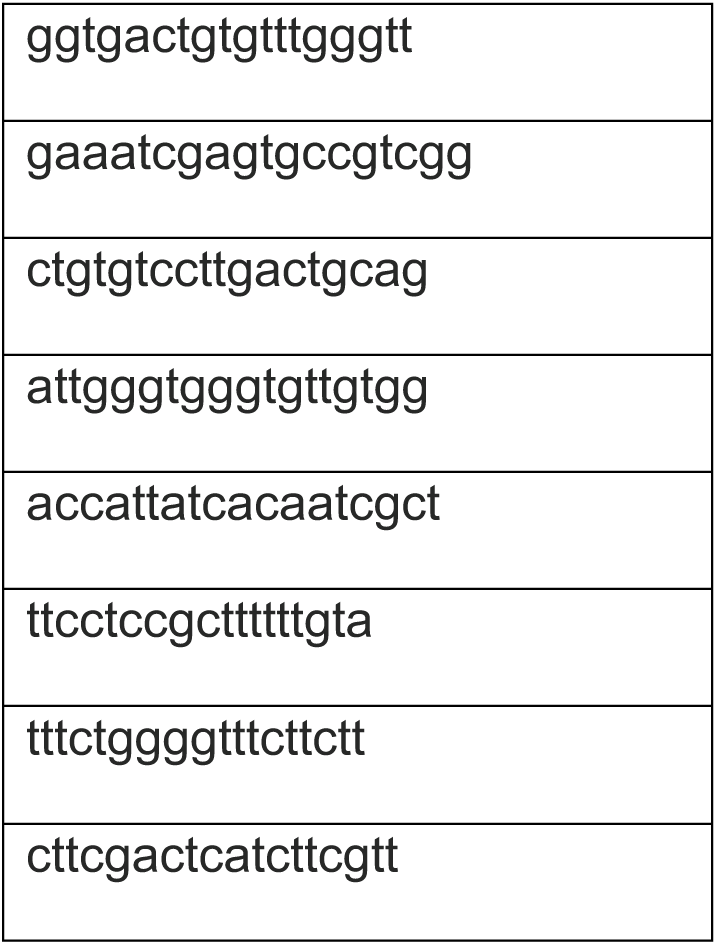

List of Hid mRNA specific probe sequences (5’-3’) designed and used in this study:

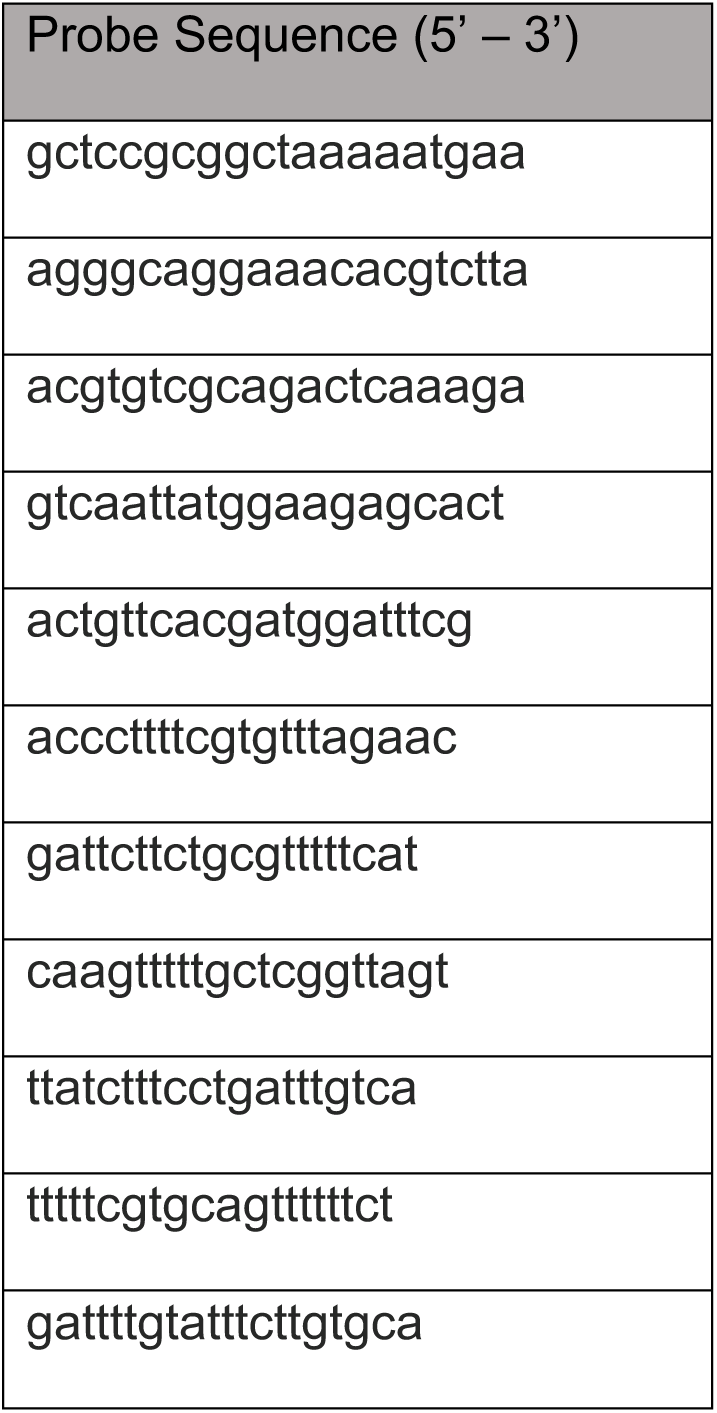
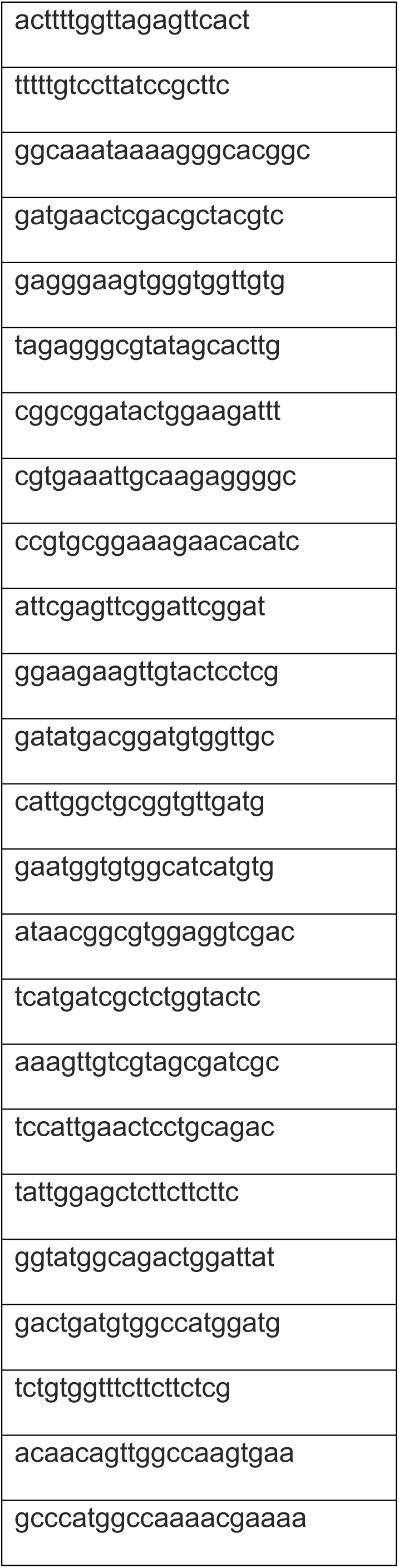
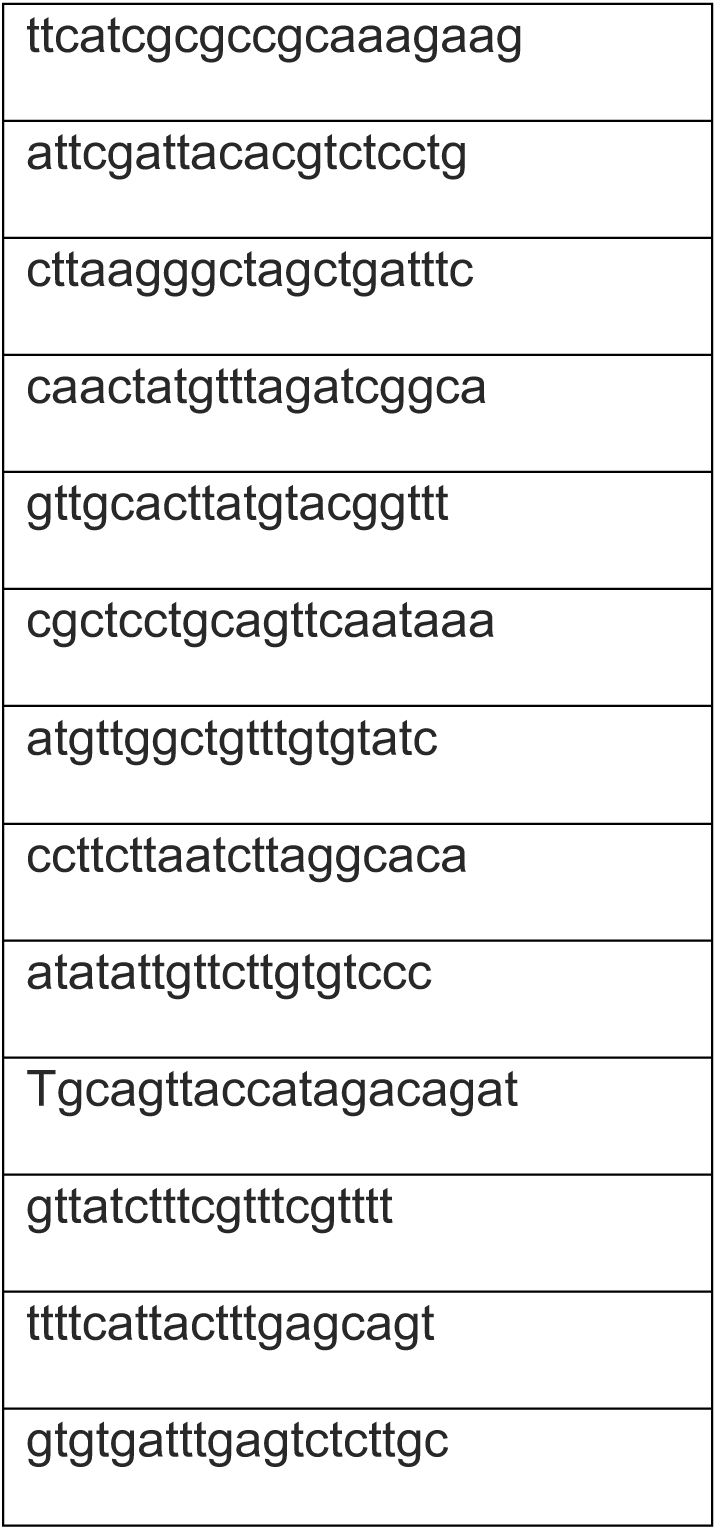

List of Sickle mRNA specific probe sequences (5’-3’) designed and used in this study:

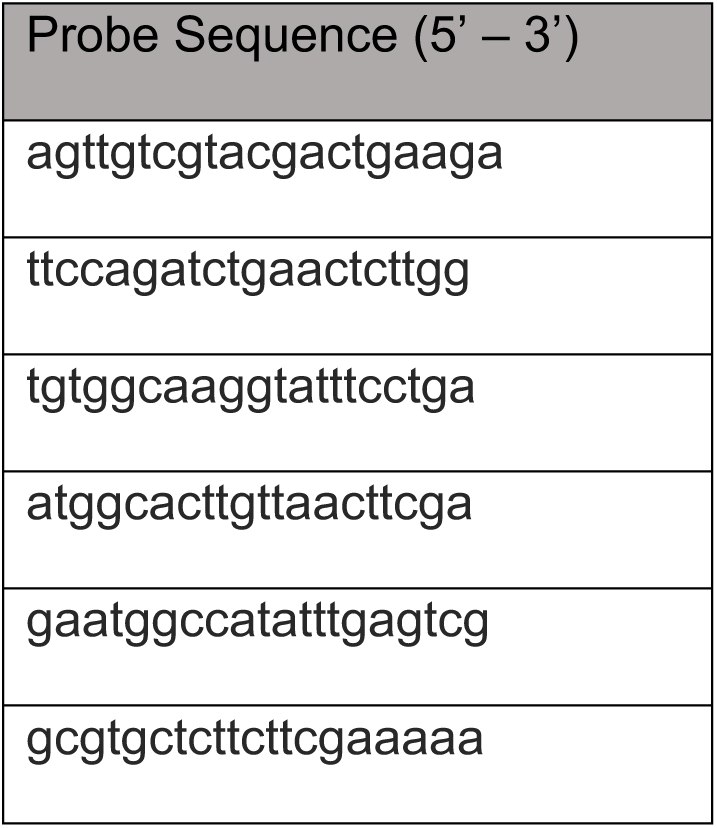
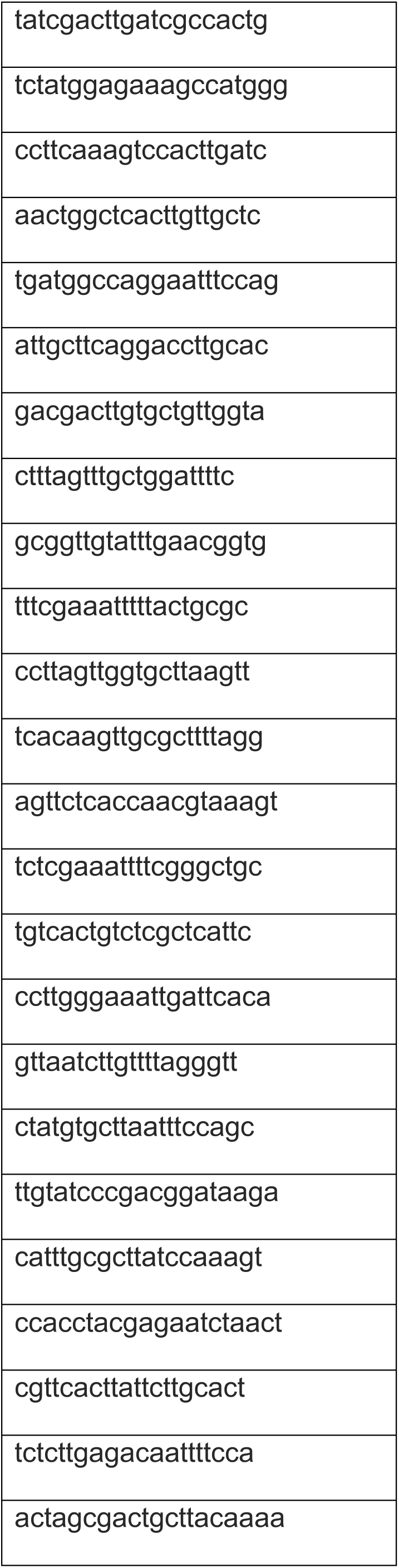
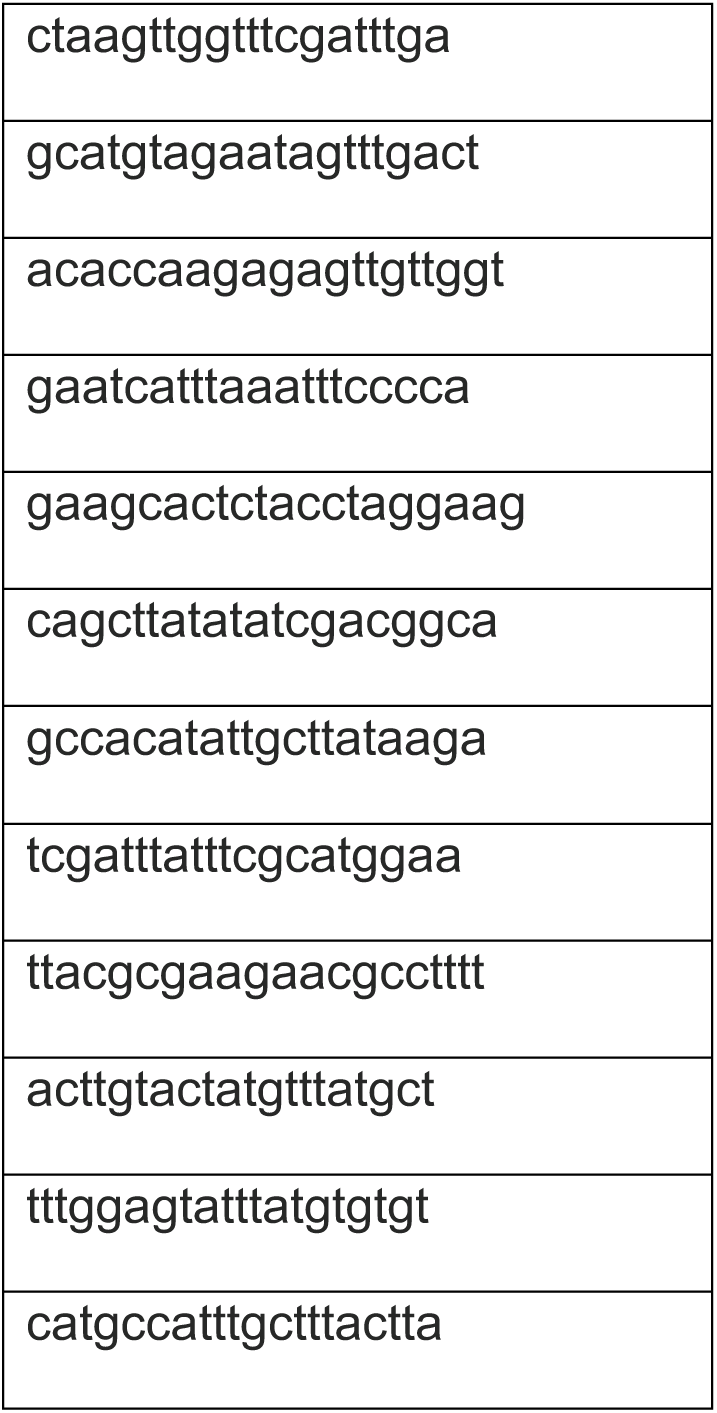

### smFISH probes used in this study not designed as part of this study

| Target | Conjugated Dye | Labels | Source |
| --- | --- | --- | --- |
| Elav | Cy5 | Pan-neuronal expression. Used to label CNS during embryonic development | Gift from L.Prieto-Godino |

## Acknowledgments

We are grateful to Troy Shirangi, Alex Gould, Kristin White and Bloomington Drosophila Stock Center (NIH P40OD018537) for generously sharing flies; Developmental Studies Hybridoma Bank (NICHD of the NIH, University of Iowa) for antibodies; Lucia Prieto-Godino for sharing Elav smFISH probe. We would like to thank Lisa Marin for her input and encouragement at the start of this project. Lucia Prieto-Godino and Richard Wingate for discussions. We also thank David Shepherd and Jon Clarke for reading drafts of the manuscript. Funding: Williams: BBSRC BB/P025552/1 and BB/L022672/1

## References

1. Simon, F., & Konstantinides, N. (2021). Single-cell transcriptomics in the Drosophila visual system: Advances and perspectives on cell identity regulation, connectivity, and neuronal diversity evolution. Developmental biology, 479, 107–122. 10.1016/j.ydbio.2021.08.001

2. Sulston, J. E., & Horvitz, H. R. (1977). Post-embryonic cell lineages of the nematode, Caenorhabditis elegans. Developmental biology, 56(1), 110–156. 10.1016/0012-1606(77)90158-0

3. Ellis, H. M., & Horvitz, H. R. (1986). Genetic control of programmed cell death in the nematode C. elegans. Cell, 44(6), 817–829. 10.1016/0092-8674(86)90004-8

4. Biehlmaier, O., Neuhauss, S. C., & Kohler, K. (2001). Onset and time course of apoptosis in the developing zebrafish retina. Cell and tissue research, 306(2), 199–207. 10.1007/s004410100447

5. Williams, J. A., Barrios, A., Gatchalian, C., Rubin, L., Wilson, S. W., & Holder, N. (2000). Programmed cell death in zebrafish rohon beard neurons is influenced by TrkC1/NT-3 signaling. Developmental biology, 226(2), 220–230. 10.1006/dbio.2000.9860

6. Lamborghini J. E. (1987). Disappearance of Rohon-Beard neurons from the spinal cord of larval Xenopus laevis. The Journal of comparative neurology, 264(1), 47–55. 10.1002/cne.902640105

7. Lamb, A. H., Ferns, M. J., & Klose, K. (1989). Peripheral competition in the control of sensory neuron numbers in Xenopus frogs reared with a single bilaterally innervated hindlimb. Brain research. Developmental brain research, 45(1), 149–153. 10.1016/0165-3806(89)90016-3

8. Yeo, W., & Gautier, J. (2003). A role for programmed cell death during early neurogenesis in xenopus. Developmental biology, 260(1), 31–45. 10.1016/s0012-1606(03)00222-7

9. Homma, S., Yaginuma, H., & Oppenheim, R. W. (1994). Programmed cell death during the earliest stages of spinal cord development in the chick embryo: a possible means of early phenotypic selection. The Journal of comparative neurology, 345(3), 377–395. 10.1002/cne.903450305

10. Hamburger, V., & Levi-Montalcini, R. (1949). Proliferation, differentiation and degeneration in the spinal ganglia of the chick embryo under normal and experimental conditions. The Journal of experimental zoology, 111(3), 457–501. 10.1002/jez.1401110308

11. Oppenheim R. W. (1991). Cell death during development of the nervous system. Annual review of neuroscience, 14, 453–501. 10.1146/annurev.ne.14.030191.002321

12. Ikonomidou, C., Bosch, F., Miksa, M., Bittigau, P., Vöckler, J., Dikranian, K., Tenkova, T. I., Stefovska, V., Turski, L., & Olney, J. W. (1999). Blockade of NMDA receptors and apoptotic neurodegeneration in the developing brain. Science (New York, N.Y.), 283(5398), 70–74. 10.1126/science.283.5398.70

13. Blanquie, O., Yang, J. W., Kilb, W., Sharopov, S., Sinning, A., & Luhmann, H. J. (2017). Electrical activity controls area-specific expression of neuronal apoptosis in the mouse developing cerebral cortex. eLife, 6, e27696. 10.7554/eLife.27696

14. Wong, F. K., Bercsenyi, K., Sreenivasan, V., Portalés, A., Fernández-Otero, M., & Marín, O. (2018). Pyramidal cell regulation of interneuron survival sculpts cortical networks. Nature, 557(7707), 668–673. 10.1038/s41586-018-0139-6

15. Truman, J. W., Moats, W., Altman, J., Marin, E. C., & Williams, D. W. (2010). Role of Notch signaling in establishing the hemilineages of secondary neurons in Drosophila melanogaster. Development (Cambridge, England), 137(1), 53–61. 10.1242/dev.041749

16. Lin, S., Lai, S. L., Yu, H. H., Chihara, T., Luo, L., & Lee, T. (2010). Lineage-specific effects of Notch/Numb signaling in post-embryonic development of the Drosophila brain. Development (Cambridge, England), 137(1), 43–51. 10.1242/dev.041699

17. Bertet, C., Li, X., Erclik, T., Cavey, M., Wells, B., & Desplan, C. (2014). Temporal patterning of neuroblasts controls Notch-mediated cell survival through regulation of Hid or Reaper. Cell, 158(5), 1173–1186. 10.1016/j.cell.2014.07.045

18. Lovick, J. K., Kong, A., Omoto, J. J., Ngo, K. T., Younossi-Hartenstein, A., & Hartenstein, V. (2016). Patterns of growth and tract formation during the early development of secondary lineages in the Drosophila larval brain. Developmental neurobiology, 76(4), 434–451. 10.1002/dneu.22325

19. Pop, S., Chen, C. L., Sproston, C. J., Kondo, S., Ramdya, P., & Williams, D. W. (2020). Extensive and diverse patterns of cell death sculpt neural networks in insects. eLife, 9, e59566. 10.7554/eLife.59566

20. Spana, E. P., & Doe, C. Q. (1996). Numb antagonizes Notch signaling to specify sibling neuron cell fates. Neuron, 17(1), 21–26. 10.1016/s0896-6273(00)80277-9

21. Lundell, M. J., Lee, H. K., Pérez, E., & Chadwell, L. (2003). The regulation of apoptosis by Numb/Notch signaling in the serotonin lineage of Drosophila. *Development (Cambridge*, England), 130(17), 4109–4121. 10.1242/dev.00593

22. Lacin, H., Chen, H. M., Long, X., Singer, R. H., Lee, T., & Truman, J. W. (2019). Neurotransmitter identity is acquired in a lineage-restricted manner in the *Drosophila* CNS. eLife, 8, e43701. 10.7554/eLife.43701

23. Harris, R. M., Pfeiffer, B. D., Rubin, G. M., & Truman, J. W. (2015). Neuron hemilineages provide the functional ground plan for the Drosophila ventral nervous system. eLife, 4, e04493. 10.7554/eLife.04493

24. Goyal, L., McCall, K., Agapite, J., Hartwieg, E., & Steller, H. (2000). Induction of apoptosis by Drosophila reaper, hid and grim through inhibition of IAP function. The EMBO journal, 19(4), 589–597. 10.1093/emboj/19.4.589

25. Lisi, S., Mazzon, I., & White, K. (2000). Diverse domains of THREAD/DIAP1 are required to inhibit apoptosis induced by REAPER and HID in Drosophila. Genetics, 154(2), 669–678. 10.1093/genetics/154.2.669

26. Yoo, S. J., Huh, J. R., Muro, I., Yu, H., Wang, L., Wang, S. L., Feldman, R. M., Clem, R. J., Müller, H. A., & Hay, B. A. (2002). Hid, Rpr and Grim negatively regulate DIAP1 levels through distinct mechanisms. Nature cell biology, 4(6), 416–424. 10.1038/ncb793

27. Miguel-Aliaga, I., & Thor, S. (2004). Segment-specific prevention of pioneer neuron apoptosis by cell-autonomous, postmitotic Hox gene activity. *Development (Cambridge*, England), 131(24), 6093–6105. 10.1242/dev.01521

28. Rogulja-Ortmann, A., Renner, S., & Technau, G. M. (2008). Antagonistic roles for Ultrabithorax and Antennapedia in regulating segment-specific apoptosis of differentiated motoneurons in the Drosophila embryonic central nervous system. *Development (Cambridge*, England), 135(20), 3435–3445. 10.1242/dev.023986

29. Peterson, C., Carney, G. E., Taylor, B. J., & White, K. (2002). reaper is required for neuroblast apoptosis during Drosophila development. Development (Cambridge, England), 129(6), 1467–1476. 10.1242/dev.129.6.1467

30. Bello, B. C., Hirth, F., & Gould, A. P. (2003). A pulse of the Drosophila Hox protein Abdominal-A schedules the end of neural proliferation via neuroblast apoptosis. Neuron, 37(2), 209–219. 10.1016/s0896-6273(02)01181-9

31. Winbush, A., & Weeks, J. C. (2011). Steroid-triggered, cell-autonomous death of a Drosophila motoneuron during metamorphosis. Neural development, 6, 15. 10.1186/1749-8104-6-15

32. Robinow, S., Talbot, W. S., Hogness, D. S., & Truman, J. W. (1993). Programmed cell death in the Drosophila CNS is ecdysone-regulated and coupled with a specific ecdysone receptor isoform. Development (Cambridge, England), 119(4), 1251–1259. 10.1242/dev.119.4.1251

33. Kimura, K. I., & Truman, J. W. (1990). Postmetamorphic cell death in the nervous and muscular systems of Drosophila melanogaster. The Journal of neuroscience : the official journal of the Society for Neuroscience, 10(2), 403–411. 10.1523/JNEUROSCI.10-02-00403.1990

34. White, K., Grether, M. E., Abrams, J. M., Young, L., Farrell, K., & Steller, H. (1994). Genetic control of programmed cell death in Drosophila. Science (New York, N.Y.), 264(5159), 677–683. 10.1126/science.8171319

35. White, K., Tahaoglu, E., & Steller, H. (1996). Cell killing by the Drosophila gene reaper. Science (New York, N.Y.), 271(5250), 805–807. 10.1126/science.271.5250.805

36. Grether, M. E., Abrams, J. M., Agapite, J., White, K., & Steller, H. (1995). The head involution defective gene of Drosophila melanogaster functions in programmed cell death. Genes & development, 9(14), 1694–1708. 10.1101/gad.9.14.1694

37. Chen, P., Nordstrom, W., Gish, B., & Abrams, J. M. (1996). grim, a novel cell death gene in Drosophila. Genes & development, 10(14), 1773–1782. 10.1101/gad.10.14.1773

38. Wing, J. P., Karres, J. S., Ogdahl, J. L., Zhou, L., Schwartz, L. M., & Nambu, J. R. (2002). Drosophila sickle is a novel grim-reaper cell death activator. Current biology : CB, 12(2), 131–135. 10.1016/s0960-9822(01)00664-9

39. Christich, A., Kauppila, S., Chen, P., Sogame, N., Ho, S. I., & Abrams, J. M. (2002). The damage-responsive Drosophila gene sickle encodes a novel IAP binding protein similar to but distinct from reaper, grim, and hid. Current biology : CB, 12(2), 137–140. 10.1016/s0960-9822(01)00658-3

40. Srinivasula, S. M., Datta, P., Kobayashi, M., Wu, J. W., Fujioka, M., Hegde, R., Zhang, Z., Mukattash, R., Fernandes-Alnemri, T., Shi, Y., Jaynes, J. B., & Alnemri, E. S. (2002). sickle, a novel Drosophila death gene in the reaper/hid/grim region, encodes an IAP-inhibitory protein. Current biology : CB, 12(2), 125–130. 10.1016/s0960-9822(01)00657-1

41. Huh, J. R., Vernooy, S. Y., Yu, H., Yan, N., Shi, Y., Guo, M., & Hay, B. A. (2004). Multiple apoptotic caspase cascades are required in nonapoptotic roles for Drosophila spermatid individualization. PLoS biology, 2(1), E15. 10.1371/journal.pbio.0020015

42. Zhou L. (2005). The ‘unique key’ feature of the Iap-binding motifs in RHG proteins. Cell death and differentiation, 12(8), 1148–1151. 10.1038/sj.cdd.4401637

43. Taghert, P. H., Doe, C. Q., & Goodman, C. S. (1984). Cell determination and regulation during development of neuroblasts and neurones in grasshopper embryo. Nature, 307(5947), 163–165. 10.1038/307163a0

44. Isshiki, T., Pearson, B., Holbrook, S., & Doe, C. Q. (2001). Drosophila neuroblasts sequentially express transcription factors which specify the temporal identity of their neuronal progeny. Cell, 106(4), 511–521. 10.1016/s0092-8674(01)00465-2

45. Mellert, D. J., Williamson, W. R., Shirangi, T. R., Card, G. M., & Truman, J. W. (2016). Genetic and Environmental Control of Neurodevelopmental Robustness in Drosophila. PloS one, 11(5), e0155957. 10.1371/journal.pone.0155957

46. Marin, E. C., Dry, K. E., Alaimo, D. R., Rudd, K. T., Cillo, A. R., Clenshaw, M. E., Negre, N., White, K. P., & Truman, J. W. (2012). Ultrabithorax confers spatial identity in a context-specific manner in the Drosophila postembryonic ventral nervous system. Neural development, 7, 31. 10.1186/1749-8104-7-31

47. Manoli, D. S., Foss, M., Villella, A., Taylor, B. J., Hall, J. C., & Baker, B. S. (2005). Male-specific fruitless specifies the neural substrates of Drosophila courtship behaviour. Nature, 436(7049), 395–400. 10.1038/nature03859

48. Shepherd, D., Harris, R., Williams, D. W., & Truman, J. W. (2016). Postembryonic lineages of the Drosophila ventral nervous system: Neuroglian expression reveals the adult hemilineage associated fiber tracts in the adult thoracic neuromeres. The Journal of comparative neurology, 524(13), 2677– 2695. 10.1002/cne.23988

49. Lee, G., Hall, J. C., & Park, J. H. (2002). Doublesex gene expression in the central nervous system of Drosophila melanogaster. Journal of neurogenetics, 16(4), 229–248. 10.1080/01677060216292

50. Rideout, E. J., Dornan, A. J., Neville, M. C., Eadie, S., & Goodwin, S. F. (2010). Control of sexual differentiation and behavior by the doublesex gene in Drosophila melanogaster. Nature neuroscience, 13(4), 458–466. 10.1038/nn.2515

51. Barolo, S., Castro, B., & Posakony, J. W. (2004). New Drosophila transgenic reporters: insulated P-element vectors expressing fast-maturing RFP. BioTechniques, 36(3), 436–442. 10.2144/04363ST03

52. Buss, R. R., Sun, W., & Oppenheim, R. W. (2006). Adaptive roles of programmed cell death during nervous system development. Annual review of neuroscience, 29, 1–35. 10.1146/annurev.neuro.29.051605.112800

53. Truman, J. W., Thorn, R. S., & Robinow, S. (1992). Programmed neuronal death in insect development. Journal of neurobiology, 23(9), 1295–1311. 10.1002/neu.480230917

54. Hobert O. (2016). Terminal Selectors of Neuronal Identity. Current topics in developmental biology, 116, 455–475. 10.1016/bs.ctdb.2015.12.007

55. Nordstrom, W., Chen, P., Steller, H., & Abrams, J. M. (1996). Activation of the reaper gene during ectopic cell killing in Drosophila. Developmental biology, 180(1), 213–226. 10.1006/dbio.1996.0296

56. Florentin, A., & Arama, E. (2012). Caspase levels and execution efficiencies determine the apoptotic potential of the cell. The Journal of cell biology, 196(4), 513–527. 10.1083/jcb.201107133

57. Nakajima, Y. I., & Kuranaga, E. (2017). Caspase-dependent non-apoptotic processes in development. Cell death and differentiation, 24(8), 1422–1430. 10.1038/cdd.2017.36

58. Tan, Y., Yamada-Mabuchi, M., Arya, R., St Pierre, S., Tang, W., Tosa, M., Brachmann, C., & White, K. (2011). Coordinated expression of cell death genes regulates neuroblast apoptosis. *Development (Cambridge*, England), 138(11), 2197–2206. 10.1242/dev.058826

59. Khandelwal, R., Sipani, R., Govinda Rajan, S., Kumar, R., & Joshi, R. (2017). Combinatorial action of Grainyhead, Extradenticle and Notch in regulating Hox mediated apoptosis in Drosophila larval CNS. PLoS genetics, 13(10), e1007043. 10.1371/journal.pgen.1007043

60. Arya, R., Sarkissian, T., Tan, Y., & White, K. (2015). Neural stem cell progeny regulate stem cell death in a Notch and Hox dependent manner. Cell death and differentiation, 22(8), 1378–1387. 10.1038/cdd.2014.235

61. Lacin, H., Zhu, Y., Wilson, B. A., & Skeath, J. B. (2014). Transcription factor expression uniquely identifies most postembryonic neuronal lineages in the Drosophila thoracic central nervous system. *Development (Cambridge*, England), 141(5), 1011–1021. 10.1242/dev.102178

62. Lacin, H., & Truman, J. W. (2016). Lineage mapping identifies molecular and architectural similarities between the larval and adult Drosophila central nervous system. eLife, 5, e13399. 10.7554/eLife.13399

63. Allen, A. M., Neville, M. C., Birtles, S., Croset, V., Treiber, C. D., Waddell, S., & Goodwin, S. F. (2020). A single-cell transcriptomic atlas of the adult *Drosophila* ventral nerve cord. eLife, 9, e54074. 10.7554/eLife.54074

64. Pollington, H. Q., Seroka, A. Q., & Doe, C. Q. (2023). From temporal patterning to neuronal connectivity in Drosophila type I neuroblast lineages. Seminars in cell & developmental biology, 142, 4–12. 10.1016/j.semcdb.2022.05.022

65. Shirangi, T. R., Wong, A. M., Truman, J. W., & Stern, D. L. (2016). Doublesex Regulates the Connectivity of a Neural Circuit Controlling Drosophila Male Courtship Song. Developmental cell, 37(6), 533–544. 10.1016/j.devcel.2016.05.012

66. Kimura, K., Ote, M., Tazawa, T., & Yamamoto, D. (2005). Fruitless specifies sexually dimorphic neural circuitry in the Drosophila brain. Nature, 438(7065), 229–233. 10.1038/nature04229

67. Ghosh, N., Bakshi, A., Khandelwal, R., Rajan, S. G., & Joshi, R. (2019). The Hox gene *Abdominal-B* uses Doublesex^F^ as a cofactor to promote neuroblast apoptosis in the *Drosophila* central nervous system. *Development (Cambridge*, England), 146(16), dev175158. 10.1242/dev.175158

68. Truman, J. W., & Ball, E. E. (1998). Patterns of embryonic neurogenesis in a primitive wingless insect, the silverfish, Ctenolepisma longicaudata: comparison with those seen in flying insects. Development genes and evolution, 208(7), 357–368. 10.1007/s004270050192

69. Biffar, L., & Stollewerk, A. (2014). Conservation and evolutionary modifications of neuroblast expression patterns in insects. Developmental biology, 388(1), 103–116. 10.1016/j.ydbio.2014.01.028

70. Prieto-Godino, L. L., Silbering, A. F., Khallaf, M. A., Cruchet, S., Bojkowska, K., Pradervand, S., Hansson, B. S., Knaden, M., & Benton, R. (2020). Functional integration of “undead” neurons in the olfactory system. Science advances, 6(11), eaaz7238. 10.1126/sciadv.aaz7238

